# Distinct genetic pathways define pre-leukemic and compensatory clonal hematopoiesis in Shwachman-Diamond syndrome

**DOI:** 10.1101/2020.06.04.134692

**Authors:** Alyssa L. Kennedy, Kasiani C. Myers, James Bowman, Christopher J. Gibson, Nicholas D. Camarda, Elissa Furutani, Gwen M. Muscato, Robert H. Klein, Kaitlyn Ballotti, Shanshan Liu, Chad E. Harris, Ashley Galvin, Maggie Malsch, David Dale, John M. Gansner, Taizo A. Nakano, Alison Bertuch, Adrianna Vlachos, Jeffrey M. Lipton, Paul Castillo, James Connelly, Jane Churpek, John R. Edward, Nobuko Hijiya, Richard H. Ho, Inga Hofmann, James N. Huang, Siobán Keel, Adam Lamble, Bonnie W. Lau, Maxim Norkin, Elliot Stieglitz, Wendy Stock, Kelly Walkovich, Steffen Boettcher, Christian Brendel, Mark D. Fleming, Stella M. Davies, Edie A. Weller, Christopher Bahl, Scott L. Carter, Akiko Shimamura, R. Coleman Lindsley

## Abstract

Shwachman-Diamond syndrome (SDS) is an inherited bone marrow failure syndrome with predisposition to developing leukemia. We found that multiple independent somatic hematopoietic clones arise early in life, most commonly harboring heterozygous mutations in *EIF6* or *TP53*. *EIF6* mutations cause functional compensation for the germline deficiency by alleviating the SDS ribosome joining defect, improving translation, and reducing p53 activation. *TP53* mutations decrease checkpoint activation without affecting ribosome assembly. We link development of leukemia with acquisition of biallelic *TP53* alterations. Our results define distinct pathways of clonal selection driven by germline fitness constraint and provide a mechanistic framework for clinical surveillance.

---

Genetic predisposition to myeloid malignancy comprises a separate disease entity in the WHO classification^1^. Diagnosis of leukemia predisposition provides potential opportunities for early intervention, but precision medicine approaches to clinical surveillance are lacking.

Shwachman-Diamond syndrome (SDS) is a genetic disorder associated with a high risk of developing myeloid neoplasms early in life^2–4^. SDS is predominantly caused by biallelic germline mutations in the *SBDS* gene^5^. The SBDS protein promotes formation of the mature, translationally active 80S ribosome by cooperating with the GTPase EFL1 to catalyze the removal of EIF6 from the 60S ribosomal subunit. In the absence of SBDS, EIF6 remains bound to the 60S subunit and sterically inhibits its joining to the 40S subunit^6^. In SDS cells, SBDS deficiency impairs eviction of EIF6 from the nascent 60S subunit, resulting in decreased ribosomal subunit joining and reduced translation efficiency^6^. Activation of cellular senescence pathways by ribosome stress incurs a global fitness defect in hematopoietic stem and progenitor cells which manifests clinically as bone marrow failure^7–9^.

Survival of patients with SDS who develop myelodysplastic syndrome (MDS) or acute myeloid leukemia (AML) is poor^10^. Therefore, a central goal in clinical care of SDS patients is to identify incipient leukemic transformation and initiate pre-emptive treatment with allogeneic stem cell transplantation. Current surveillance strategies for patients with SDS and other leukemia predisposition syndromes rely on monitoring hematologic status by serial peripheral blood counts to identify worsening cytopenias and bone marrow examinations to identify morphologic changes or development of clonal chromosomal abnormalities^11^. These tests are insensitive and detect abnormalities that are late signs of impending transformation.

The p53 tumor suppressor pathway is activated by defective ribosome biogenesis and aberrant protein translation^7,12^. Somatic *TP53* mutations have been observed in patients with SDS who develop MDS^13^, raising the possibility that next-generation sequencing could be integrated into surveillance for somatic clones with enhanced leukemia potential. However, *TP53* mutations have also been identified in SDS patients without myeloid neoplasms (MN)^14^, suggesting that additional factors must be uncovered before implementing molecular surveillance as a predictive tool in SDS. To understand the molecular pathogenesis of MN in patients with SDS, we characterized the presence and dynamics of somatic mutations in serial, clinically annotated samples collected prospectively from patients enrolled in the North American SDS Registry and studied the functional consequences of recurrently mutated pathways. Our results show that SDS patients develop frequent somatic hematopoietic clones that either bypass or compensate for the germline defect in ribosome function, and that detection of biallelic *TP53* alterations may reflect impending leukemia.

## Results

### Genetic pathways of somatic clonal expansion in SDS

We investigated genetic pathways that drive somatic hematopoietic clonal expansion and leukemogenesis in a cohort of 110 patients with a clinical diagnosis of SDS (Figure 1A). The clinical characteristics of the cohort are described in Table 1. We first used whole exome sequencing to identify somatic mutations in bone marrow aspirate samples and paired fibroblasts from 29 SDS patients (Figure 1A). All 12 patients with MN had somatic alterations also seen in sporadic MN, including point mutations in *TP53, RUNX1*, *SETBP1, BRAF, NRAS*, and *ETNK1*, or recurrent structural alterations involving chromosomes 3, 5, 7, and 20. As expected, we observed frequent interstitial deletions of chromosome 20q (8 of 17, 47%) in patients without MN (Supplementary Figure 1)^15^. Among these patients without MN, we further identified novel recurrent mutations in *EIF6* (5 of 17, 29%), suggesting that disruption of normal 60S:EIF6 function may drive clonal expansion in SDS cells.

**Table 1:** Patient Characteristics.

|  | **SDS**<br>(*n*=99) | **SDS-like**<br>(*n*=11) |
| --- | --- | --- |
| Age, median (range), years | 10.8 (0.3-49.3) | 15.6 (2.0-22.3) |
| Sex, n (%) |  |  |
| Male | 61 (61.6) | 10 (90.9) |
| Female | 38 (38.4) | 1 (9.1) |
| SDS germline mutation, n (%) |  |  |
| SBDS | 98 (98.9) | 0 (0) |
| *DNAJC21* | 1 (1.0) |  |
| Myeloid neoplasm, n (%) |  |  |
| AML | 8 (8.1) | 0 (0) |
| MDS | 7 (7.1) | 0 (0) |
| none | 84 (84.8) | 11 (100) |
| Severe bone marrow failure, n (%) |  |  |
| Yes | 6 (6.1) | 1 (9.1) |
| No | 93 (93.9) | 10 (90.9) |
| Granulocyte-colony-stimulating-factor, n (%) |  |  |
| Yes | 37 (37.4) | 0 (0) |
| No | 55 (55.6) | 1 (9.1) |
| Not Available | 7 (7.1) | 10 (90.9) |

**Figure 1:**
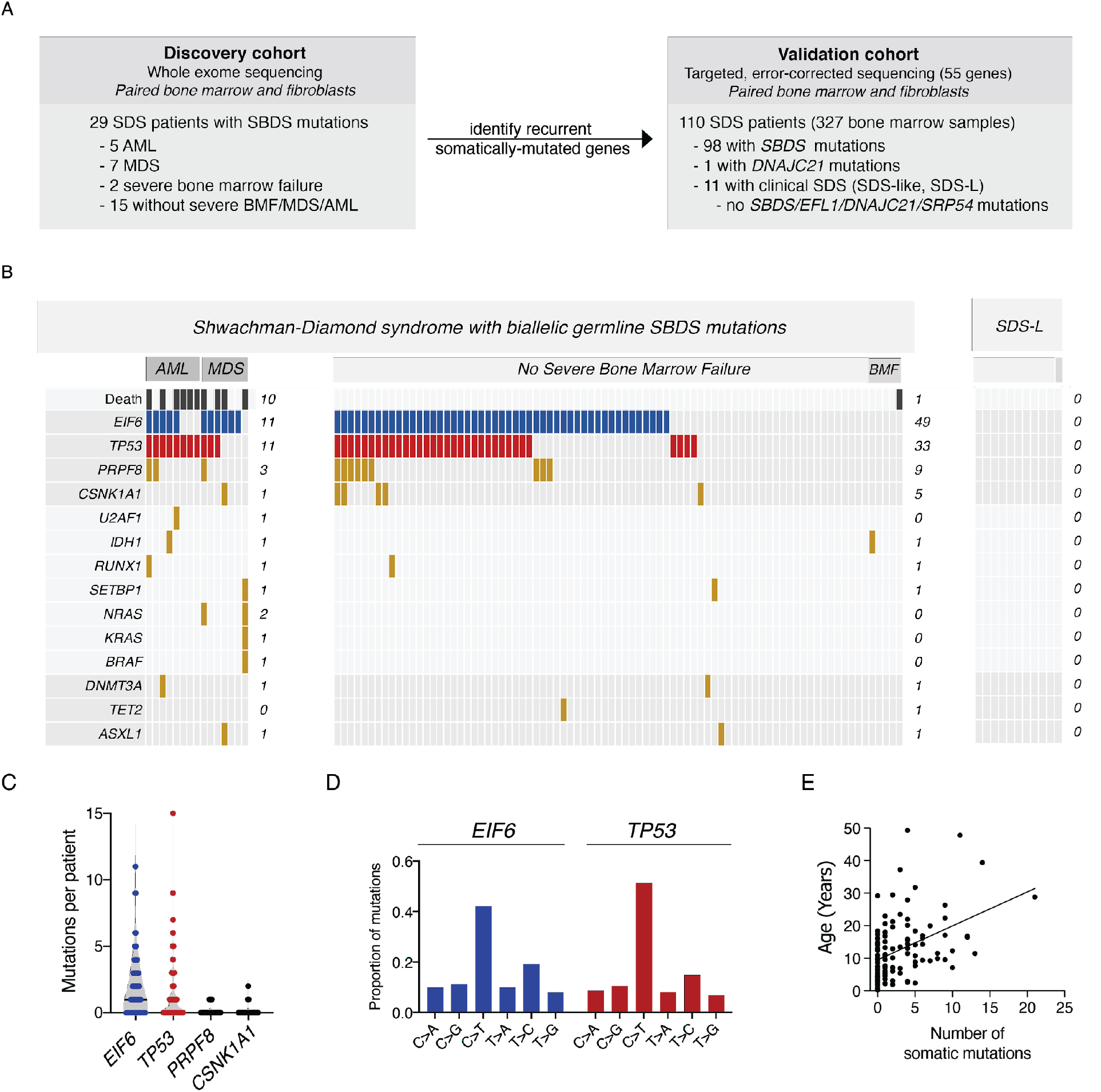
Clinical factors associated with CH in SDS patients. **A,** Schema of genomic analysis **B,** A co-mutation plot showing somatic mutations in individual genes as labeled on the left. Mutations are depicted by colored bars and each column represents an individual patient in the indicated study cohort. The sum total of each event or mutation are tabulated to the right of each plot. **C,** Number of mutations per patient in each of the four most frequently mutated genes:*TP53*, *EIF6*, *PRPF8* and *CSNK1A*. **D,** Base pair substitutions of somatic mutations in *TP53* and *EIF6*. **E,** Total number of somatic mutations by age in patients with biallelic germline *SBDS* mutations, based on targeted deep sequencing.

We next performed targeted validation of candidate gene mutations in paired bone marrow and fibroblast samples from the whole cohort. We sequenced 55 genes, including those recurrently mutated in the discovery exome cohort as well as genes associated with sporadic MN (Supplementary Table 1). To detect clones present at low abundance [0.1% variant allele fraction (VAF)], we used a platform that incorporated duplex unique molecular identifiers, thereby enabling computational suppression of sequencing artifacts.

We initially focused our analysis on the most recent sample from each patient. We detected 327 somatic mutations in 74 of 98 (76%) SDS patients with germline *SBDS* mutations (median 2 mutations/patient, range 0-21), and no mutations in patients with SDS-like (SDS-L) disease, which are patients with some clinical features of SDS without disease defining mutations (*SBDS*, *EFL1*, *DNAJC21*, *SRP54*). The most frequently mutated genes were *EIF6 (*60 of 98, 61%), *TP53 (*44 of 98, 45%), *PRPF8* (12 of 98, 12%), and *CSNK1A1* (6 of 98, 6%)(Figure 1B). Secondary somatic *SBDS* mutations were found in 3 patients and no other genes were mutated in more than two patients. Among 74 patients with somatic mutations, 52 (72.2%) had multiple mutations, frequently affecting the same gene (Figure 1C). The most common base substitution in somatic *EIF6* and *TP53* variants was a cytosine-to-thymine (C→T) transition (Figure 1D), which is the predominant mutational signature associated with normal aging hematopoietic stem cells (HSCs), sporadic clonal hematopoiesis (CH), and AML^16–19^.

### Clinical factors associated with somatic mutations

*TP53* mutations were more common in SDS patients with germline *SBDS* mutations and MN than in those without MN (73.3% versus 39.8%, p=0.023), while *EIF6*, *CSNK1A1*, and *PRPF8* were not associated with MN. In univariate analysis, the presence of any somatic mutation was associated with older age (median 12.9 versus 4.7 years, p=0.0001), as were mutations in individual genes (*TP53,* p=0.0002; *EIF6,* p=0.0042; *PRPF8,* p=0.0461)(Figure 1E). Logistic regression adjusting for age, sex, and the presence of MN showed that age was independently associated with the presence of any somatic mutation (OR=1.1, for each one-year increase in age, 95% CI 1.1-1.2, p=0.0017). Further, the total number of somatic mutations per patient was positively associated with age and MN [β(se)=0.50 (0.2071), p=0.0165] in a Poisson regression model adjusted for the same variables.

### EIF6 mutations are highly recurrent and specific to SDS

Across all samples, we identified 265 *EIF6* mutations (Figure 2A), all of which were in patients with germline *SBDS* mutations. We did not detect any *EIF6* mutations in control cohorts, including patients with SDS-L disease (n=11), patients with other leukemia predisposition disorders (germline *GATA2* deficiency syndrome, n=32; telomere biology disorders, n=5; germline *SAMD9*/*SAMD9L* mutations, n=5), or adults with sporadic AML (n=39). *EIF6* truncating mutations were distributed throughout the coding region, whereas missense mutations were predominantly located in regions encoding conserved secondary structure (Figure 2B).

**Figure 2:**
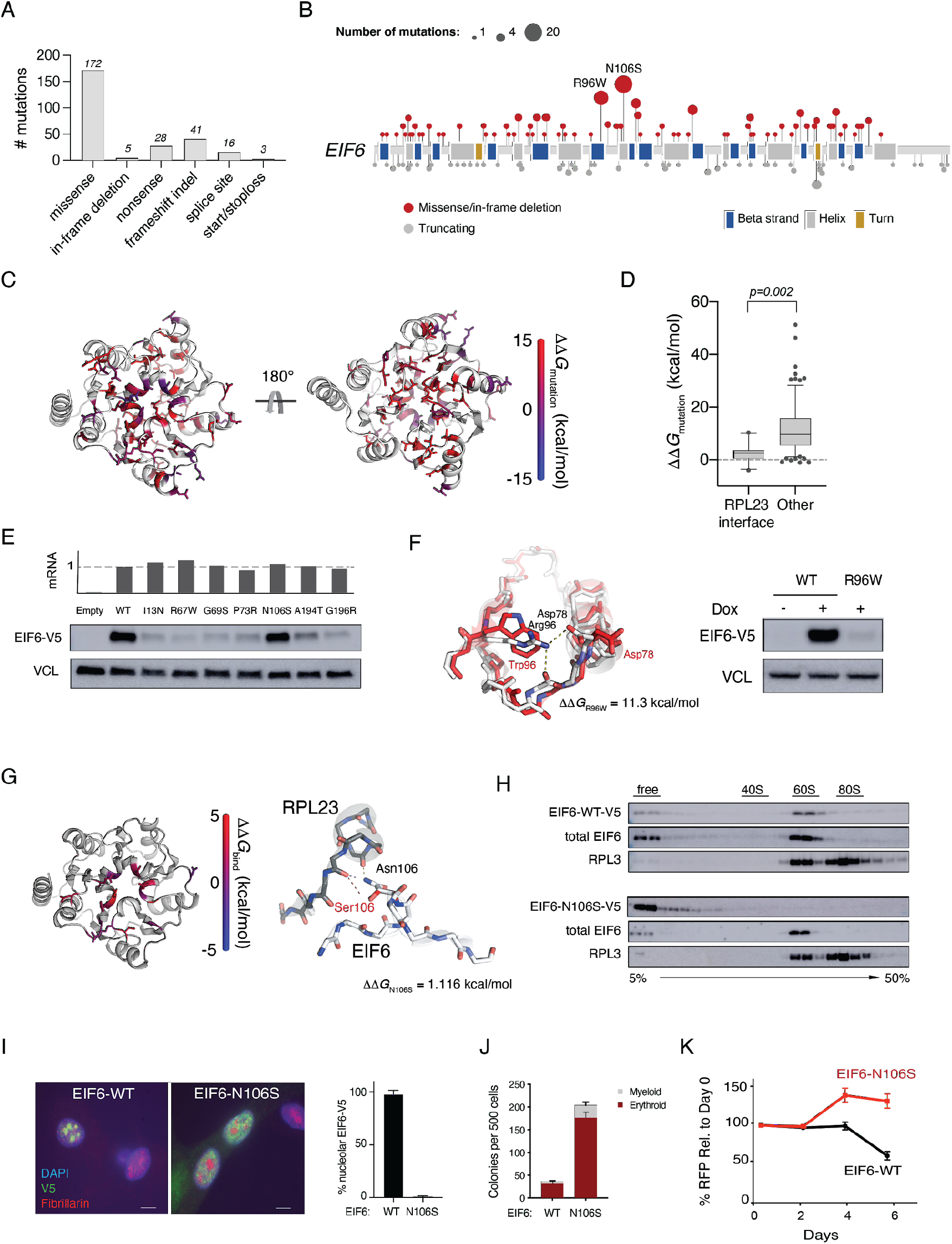
EIF6 somatic missense mutations alter EIF6 protein stability or function to improve cell fitness. **A,** Types of somatic *EIF6* mutations, **B**, Number and location of *EIF6* mutations according to variant type **C,** Impact on the calculated energy of the folded state (ΔΔ*G*_mutation_) of *EIF6* missense mutations. Mutant residue colored according to ΔΔG value. **D,**ΔΔ*G*_mutation_ of EIF6 missense mutations located at the RPL23 binding interface versus other EIF6 missense mutations located in the remainder of EIF6. **E,** Relative levels of EIF6 mRNA and V5 immunoblot from K562 cells 48 hours after doxycycline treatment. **F,** Left panel: In silico modeling of EIF6-R96W. Right panel: V5 and VCL immunoblots of K562 cells with inducible EIF6-R96W versus V5-wild type EIF6 48 hours after doxycycline treatment. **G,** Left panel: Change in the energy of binding (ΔΔ*G*_bind_) of missense mutations at RPL23 interface. Mutant residues are colored according to ΔΔ*G*_bind_. Right panel: In silico modeling of EIF6 N106S mutation. **H,** V5, EIF6, and RPL3 immunoblots of sucrose gradient fractions from polysome profiles of doxycycline treated K562 cells expressing V5-EIF6-WT or V5-EIF6-N106S. **I,** Immunofluorescence of V5-EIF-WT or V5-N106S -EIF6 protein in SDS patient-derived fibroblasts, V5 (green), fibrillarin (red), and DAPI (blue). Right panel: quantification of V5 nucleolar signal from 3 independent experiments **J.** Quantification of colony forming units from sorted CD34+ transduced with shSBDS-GFP and either EIF6-WT-RFP or EIF6-N106S-RFP plated in triplicate for 3 independent experiments. **K,** Competitive growth of shSBDS cells transduced with either EIF6-WT-RFP or EIF6-N106S-RFP after doxycycline treatment.

To study the consequences of *EIF6* missense mutations, we generated a homology model that closely matches the EIF6 structures from *Methanocaldococcus jannaschii^20^*, *Saccharomyces cerevisiae^20,21^*, and *Dictyostelium discoideum^22^* (Supplementary Figure 2). To evaluate the impact of mutations on protein stability, we modeled the effect of each mutation on the energy of the folded state of EIF6 and compared this to the wild-type protein (ΔΔ*G*_mutation_)(Figure 2C). Missense mutations located at the EIF6:RPL23 interface were not predicted to destabilize the protein (median ΔΔ*G*_mutation_=2.66 kcal/mol, 95% CI 0.29 to 3.73)(Figure 2D and Supplementary Table 2). In contrast, mutations not located at binding interfaces (median ΔΔ*G*_mutation_=9.83 kcal/mol, 95% CI 7.37 to 12.37) and those located at the non-60S interface with EFL1 (median ΔΔ*G*_mutation_=10.04 kcal/mol, 95% CI 5.79 to 24.93)(Figure 2D) were strongly destabilizing.

### EIF6 missense mutations disrupt 60S:EIF6 function by two mechanisms

We cloned patient-derived mutations with different predicted functional consequences and generated K562 leukemia cell lines that expressed wild-type or mutant EIF6 cDNA with a C-terminal V5-epitope tag under the control of a doxycycline-inducible promoter. We measured EIF6 protein levels and mRNA expression after 48 hours of doxycycline treatment and found that six mutants (I13N, R67W, G69S, P73R, A194T, G196R) had reduced levels of EIF6 protein compared with EIF6^WT^, despite comparable abundance of mutant mRNA (Figure 2E). Mutant EIF6 protein abundance was increased after treatment with the proteasome inhibitor MG-132 ( Supplementary Figure 3A). These results indicate that *EIF6* missense mutations can cause functional inactivation via protein destabilization.

The two most common recurrent mutations in the cohort were *EIF6* p.N106S and *EIF6* p.R96W, found in 20% and 13% of SDS patients, respectively. In the EIF6 homology model, R96W disrupts hydrogen bonds and is predicted to destabilize the protein (ΔΔ*G*_mutation_=11.3 kcal/mol), while N106S is predicted to be stable (ΔΔ*G*_mutation_=1.12 kcal/mol). Consistent with these models, the level of EIF6^R96W^ protein was markedly reduced compared with EIF6^WT^ (Figure 2F) and the level of EIF6^N106S^ protein was similar to EIF6^WT^ (Figure 2G).

Since N106 is highly conserved and located at the interface between EIF6 and the 60S ribosomal protein RPL23 (Figure 2G), we tested the hypothesis that N106S impairs the EIF6:60S interaction. In the homology model, N106S had significantly increased energy of EIF6:RPL23 binding (ΔΔ*G*_bind_)^23^ compared with mutations not located at the EIF6:RPL23 interface. To directly analyze the impact of N106S on this interaction, we conducted sucrose gradient polysome profiling of lysates from cells expressing V5-tagged EIF6^WT^ or EIF6^N106S^, followed by western blotting across the gradient fractions. While EIF6^WT^ was primarily present in the 60S fractions^24,25^, EIF6^N106S^ was found only in the free fractions and was absent from the 60S fractions (Figure 2H). Consistent with this result, EIF6^WT^ was distributed normally in the cytoplasm and nucleolus^21,26^ and EIF6^N106S^ was detectable only in the cytoplasm (Fig 2I).

To assess the effect of EIF6^N106S^ on the functional competency of SDS hematopoietic stem and progenitor cells, we expressed either EIF6^WT^ or EIF6^N106S^ in SBDS-deficient human CD34+ cells and quantified hematopoietic colony formation. Both myeloid and erythroid colonies were more abundant with EIF6^N106S^ compared with EIF6^WT^ (Figure 2J). Similarly, SBDS-deficient K562 leukemia cells expressing EIF6-N106S displayed increased overall growth compared to cells expressing EIF6^WT^ (Figure 2K and Supplementary Figure 3B).

### EIF6 and TP53 mutations alleviate p53 activation

SBDS deficiency impairs ribosome assembly and results in reduced abundance of the mature 80S ribosome, concomitant accumulation of free 60S ribosome subunits^27,28^, and upregulation of p53-dependent cellular stress pathways in SDS patient bone marrow^29^ and SDS mouse models^8^. Somatic mutations that reduce p53 activation could thus drive selective clonal advantage either by rescuing the underlying defect in ribosome maturation or by directly inactivating *TP53*.

To investigate the effects of *EIF6* and *TP53* mutations on ribosome maturation, protein translation, and p53 target gene activation in SBDS-deficient cells, we introduced shRNAs that targeted *EIF6* or *TP53,* or a control shRNA targeting luciferase into primary SDS patient-derived bone marrow fibroblasts (Supplementary Figure 4A). Using sucrose gradient polysome profiling, we found that knockdown of *EIF6*, but not knockdown of *TP53*, resulted in an increased ratio of 80S:60S ribosomal subunits relative to control (Figure 3A and Supplementary Figure 4B). Consistent with these distinct effects on ribosome maturation, knockdown of *EIF6*, but not knockdown of *TP53*, improved the SDS-associated impairment of protein synthesis, as measured by incorporation of O-propargyl-puromycin into nascent peptides (Figure 3B). Despite their different impact on the SDS ribosome joining defect, knockdown of either *EIF6* or *TP53* resulted in reduction of *CDKN1A* induction in SDS fibroblasts (Figure 3C). Together, these results indicate that *EIF6* and *TP53* mutations have distinct effects on ribosome joining and global protein synthesis, but share a common downstream effect of reducing p53 pathway activation.

**Figure 3:**
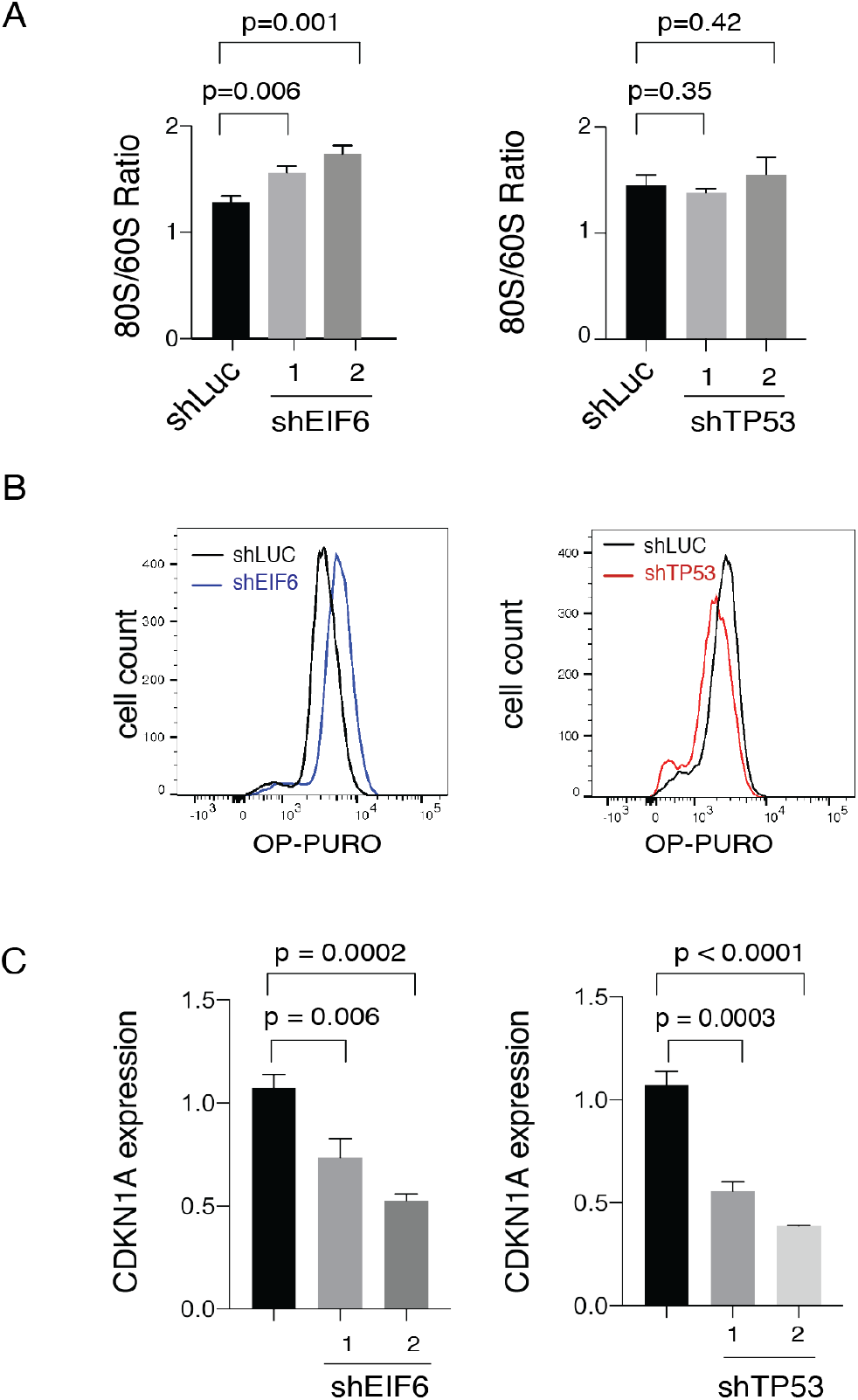
*EIF6* and *TP53* mutations attenuate p53 activation via different mechanisms. **A**, Quantification of 80S:60S ratio from polysome profiles in SDS patient-derived primary fibroblasts transduced with shRNAs targeting luciferase, EIF6 (left panel) or TP53 (right panel). **B,** OP-Puro incorporation in primary SDS patient-derived fibroblasts transduced with shRNAs targeting luciferase, EIF6 (left panel) or TP53 (right panel). **C,** Relative *CDKN1A* expression in SDS patient-derived fibroblasts transduced with either shLUC control or shEIF6 (left panel) and shTP53 (right panel).

### Independence of TP53 and EIF6 mutated clones in SDS

Among SDS patients with *TP53* mutated CH, 90.9% (30 of 33) had concurrent *EIF6* mutations, raising the possibility that *TP53* and *EIF6* mutations cooperate to drive clonal progression. To distinguish whether *TP53* and *EIF6* mutations arise in separate clones or together within the same clones, we performed single cell DNA sequencing from patients with clonal hematopoiesis who had multiple *EIF6* and *TP53* mutations detected by bulk sequencing.

Using a custom panel covering 7 genes implicated in SDS or sporadic CH and 43 single nucleotide polymorphism (SNP) loci on chromosomes 7, 17, and 20, we sequenced 33,426 cells from 6 patients with CH. The number (Figure 4A) and VAF (Figure 4B) of gene mutations detected by bulk sequencing in each patient is shown in Figure 4. Single cell genotyping was successful for 84.4% of all targeted mutations that were observed by bulk DNA sequencing and undetected mutations were restricted to low abundance clones (median VAF 0.0032, range 0.0022 to 0.0087). Using this single cell approach, we found that somatic mutations were almost always present in independent clones: among the 50 clones we identified: 24 had a sole *EIF6* mutation, 21 had a sole *TP53* mutation, and 3 had a sole *CSNK1A1* mutation (Figure 4C). One patient (SDS-026) had a clone with concurrent mutations in *TP53* and *EIF6*, where *TP53* p.R248Q defined the founding clone and *EIF6* p.S86A defined a subclone. In another patient (SDS-072), we observed a founding clone with *EIF6* p.M1T and a subclone with *TET2* p.E227* mutation.

**Figure 4:**
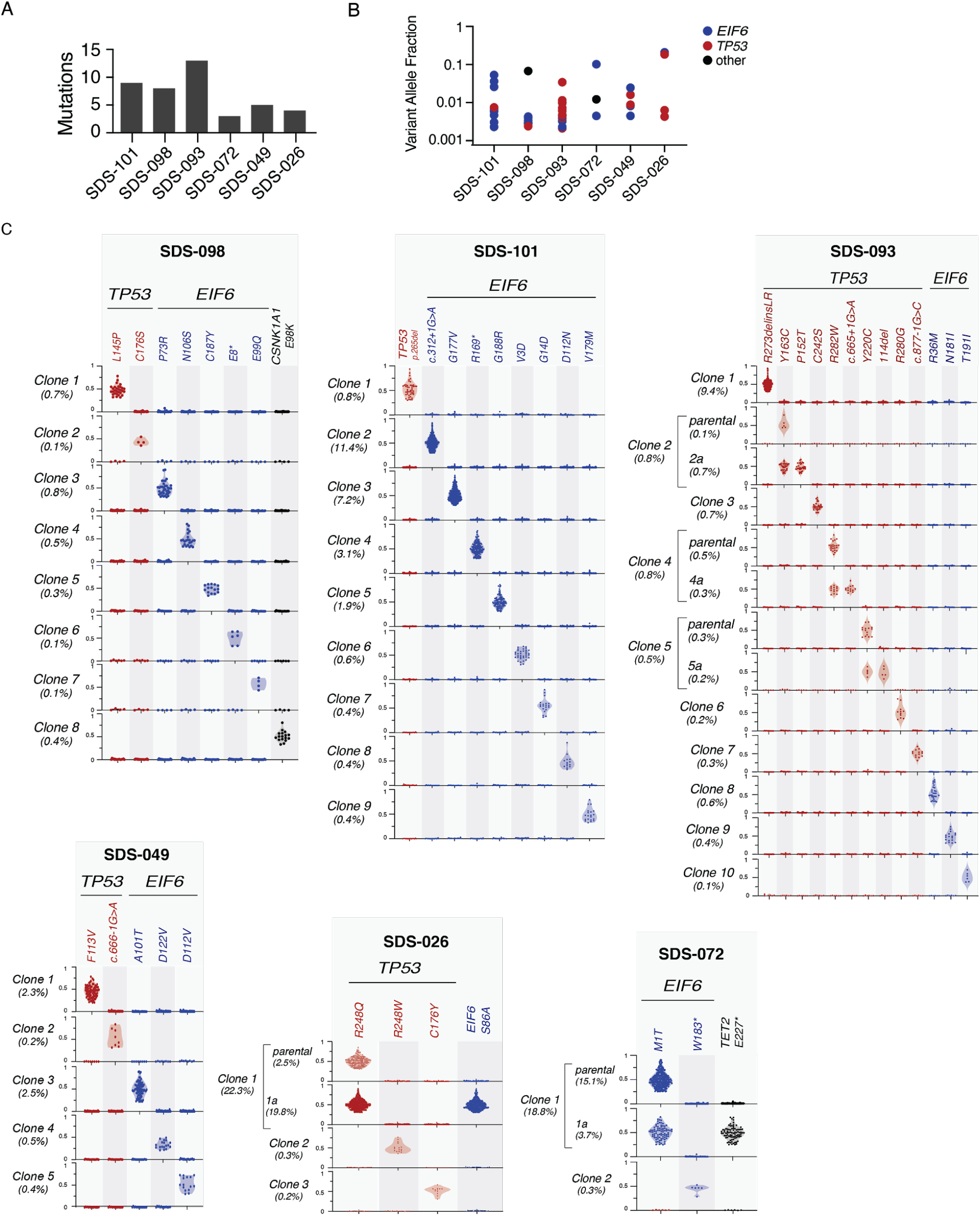
Independence of *TP53* and *EIF6* mutated clones in SDS patients. **A,** Number of somatic mutations detected in each patient by bulk DNA sequencing **B,** corresponding VAF of *TP53* (red), *EIF6* (blue) or other (black) mutation. **C,** Clonal hierarchy of mutations determined by single cell sequencing amongst six patients with SDS. Each row represents a unique clone or subclone and the frequency of each clone is indicated to the left. Columns reflect the genotype status of each mutation in each clone, and all depicted clones have complete genotyping at all loci. The y-axis indicates single cell VAF from 0 to 1, where 0 is absent, 0.5 is heterozygous mutation, and 1 is homo/hemizygous. Each dot reflects a single cell, colored according to gene mutation, *TP53* (red), *EIF6* (blue), *CSNK1A1* (black) and the frequency distribution of the data points reflected by shaded violin plots.

### Clonal hematopoiesis in SDS patients

CH in individuals without germline predisposition is associated with older age and usually involves single mutations affecting *DNMT3A*, *TET2*, or *ASXL1^19,30^*. Among 83 SDS patients without a MN diagnosis, 60 (72%) had detectable CH, 40 of whom had more than one mutation (median 3, range 1-21). Two of these patients had CH defined by clonal cytogenetic alterations in the absence of point mutations. Recurrent mutations in *EIF6* (49 of 83, 59.0%), *TP53* (33 of 83, 39.8%), *PRPF8* (9 of 83, 10.8%) and *CSNK1A1* (6 of 83, 7.2%) composed 96.9% of all somatic mutations, while typical CH mutations such as *DNMT3A*, *TET2*, or *ASXL1*, were rare (n=1 for each) (Figure 5A). CH mutations were present at low abundance irrespective of the affected gene, including *EIF6* (median VAF 0.0047, range 0.002-0.282), *TP53* (0.0044, range 0.002-0.193), *PRPF8* (0.0052, range 0.002-0.375), and *CSNK1A1* (0.0053, range 0.002-0.100) (Figure 5B). CH was detectable in 27 of 46 patients (59%) 10 years old and younger, 24 of 27 patients (89%) 11 to 20 years old (89%) and 10 of 10 patients (100%) 21 years or older (Figure 5C).

**Figure 5:**
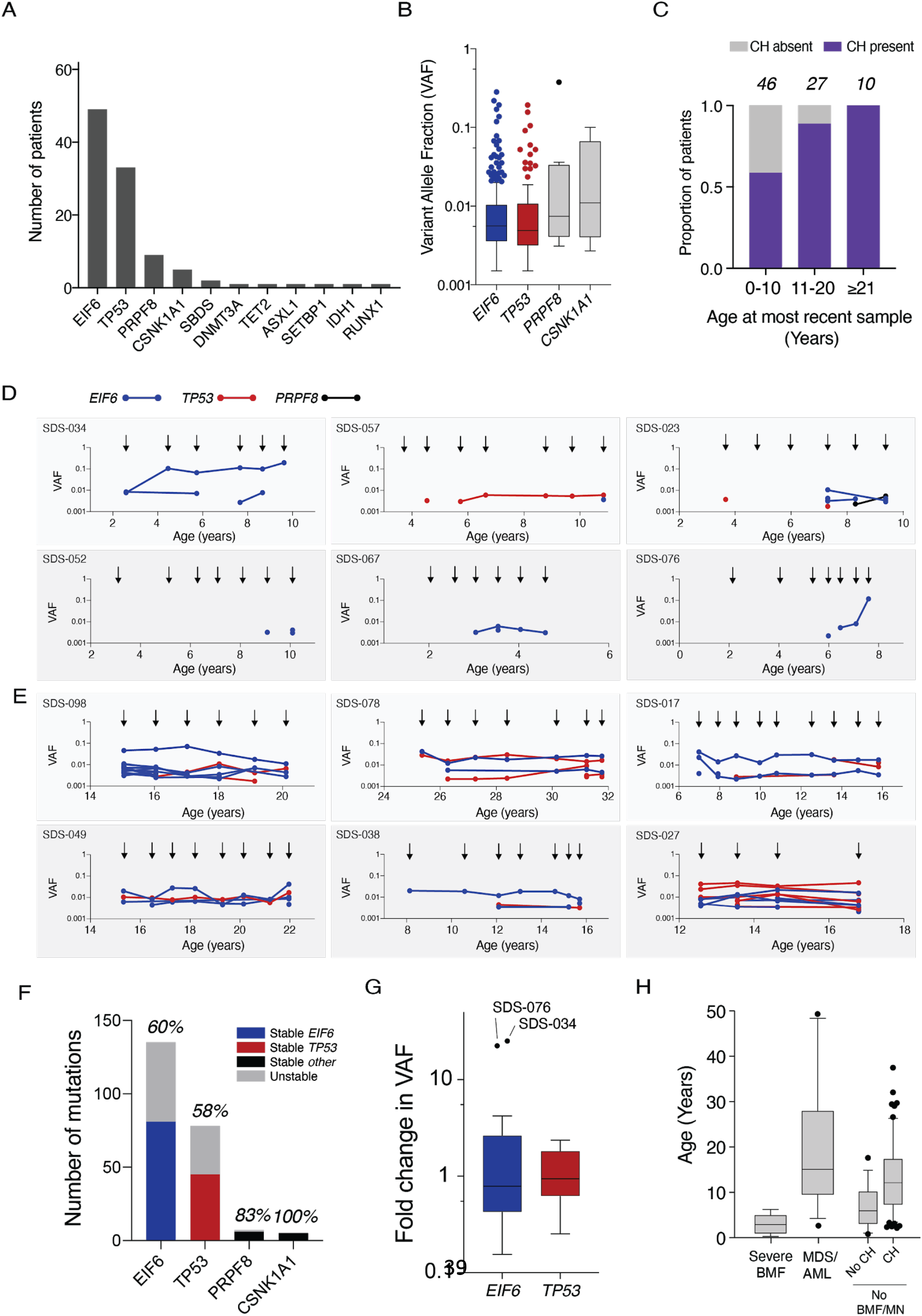
CH in SDS patients. **A**, Frequency of mutations in the indicated genes among the 58 SDS patients with CH. **B,** VAFs in the indicated genes among SDS patients with clonal hematopoiesis. Horizontal lines within boxes indicate median VAF. Boxes span the 25th and 75th percentiles, whiskers extend to the 10th and 90th percentiles, and outliers are represented by dots. **C,** Proportion of patients in the study cohort per decade of age with detectable CH. **D,E** Shown is the VAF of each somatic *EIF6* (blue), *TP53* (red) or *CSNK1A* mutation (black) from (d) six patients who developed clonal hematopoiesis in the first decade of life and (e) six patients who were found to have clonal hematopoiesis in their second or third decade of life. Arrows indicate timing of sample acquisition. Points represent the VAF for detected mutations **F,** Among patients with CH that had serial samples, the total number somatic *EIF6*, *TP53*, *PRPF8*, *and CSNK1A1* mutations and the proportion that are persistently detected across multiple samples are shown. **G,** Fold change in VAF of all somatic *EIF6* (blue) and *TP53* (red) mutations from time of first detection to time of most recent detection in patients with CH. **H,** Ages of patients at diagnosis of severe bone marrow failure (BMF), myeloid neoplasm (MN), or no BMF/MDS with or without CH at last follow up.

To assess the onset, persistence, and dynamics of CH mutations over time, we sequenced 208 serial samples from 49 SDS patients with CH (median 4 samples, range 2-11). We first analyzed serial samples from 6 patients who had developed CH prior to 10 years of age and for whom the initial sample was obtained at age 3 years or younger. In 5 cases, stable CH developed at older ages (3 to 10 years old), while in one case (SDS-034), a stable *EIF6* mutation was detected at time of first sampling at age 2 (Figure 5D). Among 6 older patients whose last sample was obtained between ages 15 to 31 years, stable CH was detectable at the earliest available time point, 5-10 years prior (Figure 5E).

Among all 49 patients with CH who had serial samples, most mutations persisted stably across multiple timepoints, including 81 of 135 *EIF6* mutations (60%), 45 of 78 *TP53* mutations (58%), 6 of 7 *PRPF8* mutations (86%), and 5 of 5 *CSNK1A1* mutations (100%)(Figure 5F). Among persistent clones, we measured the mutation allele burden across serial timepoints and found that most clones remained stable at low VAF over time, with little change in relative abundance between initial detection and the most recent sample (Figure 5G). None of the patients with CH involving *EIF6* or *TP53* had severe marrow failure despite the higher median age of the CH group compared to the group who developed severe marrow failure (Figure 5H).

### Clonal evolution and development of leukemia in SDS

The diagnosis of MN was associated with the presence of somatic *TP53* mutations. However, *TP53* mutations were common in SDS clonal hematopoiesis and typically either stable without hematologic progression across years of observation or detectable only at a single timepoint. Therefore, we sought to identify additional genetic characteristics of leukemias with somatic *TP53* mutations that distinguish CH clones with high-risk of transformation from those likely to remain clinically stable.

We analyzed exomes from seven patients with *TP53* mutated MN for allelic imbalances at the *TP53* locus by evaluating the total copy ratio (tCR) and SNP VAF across chromosome 17. We found that all seven patients had biallelic alteration of *TP53*, occurring by one of three mechanisms based on the number of *TP53* mutations (1 vs. 2 or more) and the presence of *TP53* deletion or copy-neutral loss of heterozygosity (CN-LOH). We observed 4 cases with monoallelic *TP53* mutations and 17p CN-LOH, 1 with monoallelic *TP53* mutation and 17p deletion, and 2 with biallelic *TP53* mutations (Figure 6A). In each case, the *TP53* mutations were present at high cancer cell fraction, indicating that they were likely present in all cells of the leukemic clone (Figure 6B). Among *TP53* mutated MN, 3 of 7 also harbored somatic mutations in typical myeloid drivers, including subclonal mutations in *NRAS* (n=2), *KRAS*, or *PTPN11*. Somatic mutations in genes encoding effectors of RAS/MAPK signaling (*NRAS*, *KRAS*, *PTPN11*, *CBL*, *FLT3*, *RIT1*, *KIT*) were rarely present in samples from patients without morphologic transformation.

**Figure 6:**
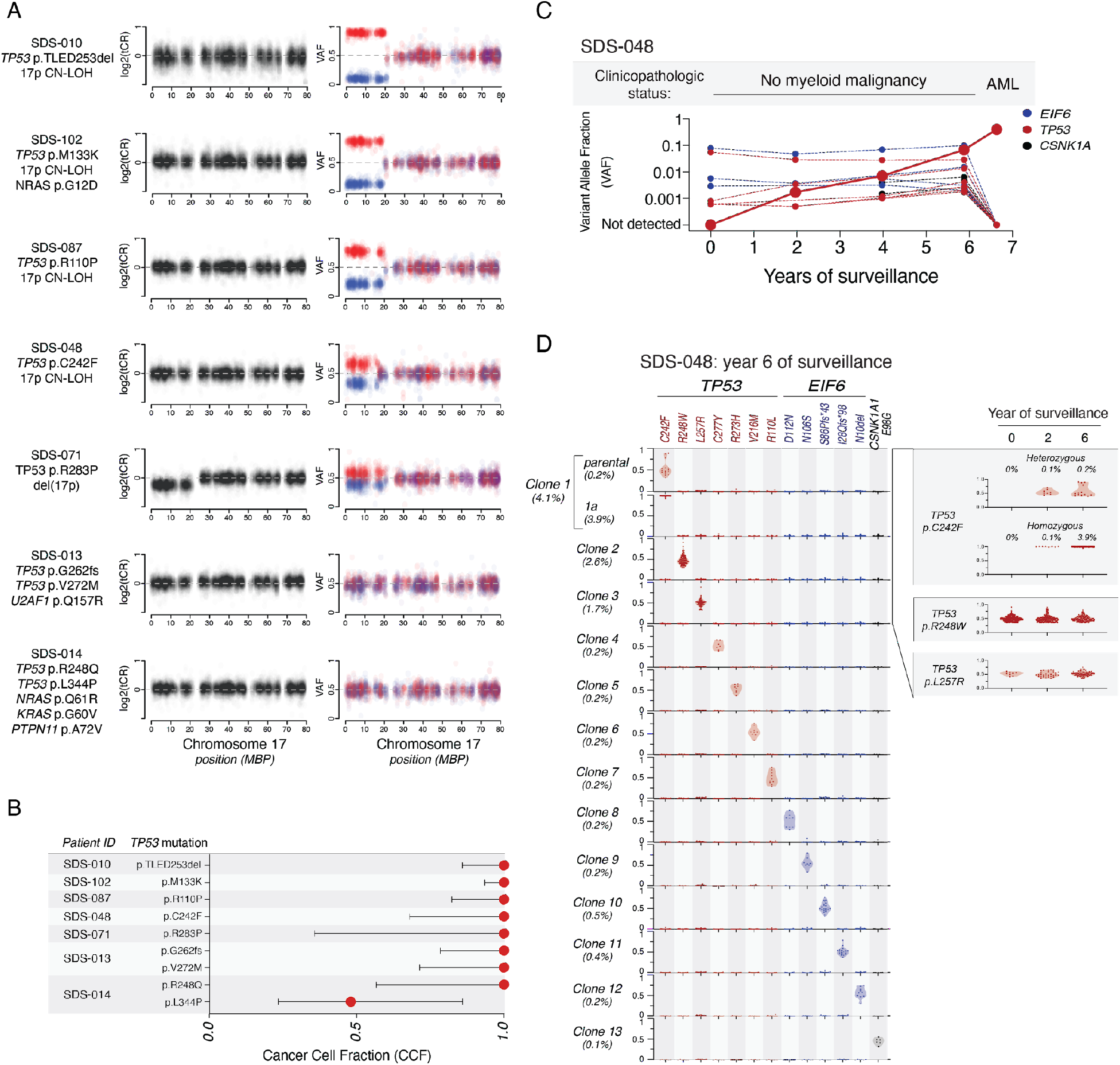
Biallelic TP53 inactivation and myeloid neoplasia in patients with SDS. **A,** Total copy ratio (tCR, denoted in black) and phased SNP-VAF (denoted in red/blue) across chromosome 17. **B,** Cancer cell fraction of somatic *TP53* mutations in samples analyzed in panel A. **C,** Shown are the clinicopathologic status and VAF of somatic *TP53* (red) *EIF6* (blue) and *CSNK1A* (black) mutations from bulk sequencing of serial samples from SDS-048, a patient with SDS who progressed to AML. **D,** Single cell sequencing demonstrating clonal hierarchy from SDS-048 at 6.5,2.5, and 0.5 years prior to development of AML. Each row represents a unique clone or subclone and the frequency of each clone is indicated to the left. Columns reflect the genotype status of each mutation in each clone, and all depicted clones have complete genotyping at all loci. The y-axis indicates single cell VAF from 0 to 1, where 0 is absent, 0.5 is heterozygous mutation, and 1 is homo/hemizygous. Each dot reflects a single cell, colored according to gene mutation, *TP53* (red), *EIF6* (blue), *CSNK1A1* (black) and the frequency distribution of the data points reflected by shaded violin plots. Shown on the right is a timecourse indicating dynamics of the pre-leukemic p.C242F mutated clone and two independent *TP53* mutated clones that did not transform.

In 4 of 15 patients with morphologically defined MN, we did not detect somatic *TP53* mutations. All four of these patients had MDS. In two, we identified mutations in canonical myeloid driver genes, including one with *SETBP1*, *BRAF*, *NRAS*, and *KRAS* mutations and one with *ASXL1* and *ETNK1* mutations. Of the remaining two patients without point mutations, one had deletion of chromosome 7q and the other was diagnosed with MDS based on morphologic dysplasia without cytogenetic abnormalities or increased blasts.

### Early identification of leukemic subclones

Early detection of leukemia-associated genetic alterations could identify clones with increased leukemic potential prior to clinical transformation. We therefore sought to define the latency between detection of these mutations and clinical progression in a patient who developed AML despite having stable blood counts and no morphologic evidence of MN on bone marrow examinations during standard clinical surveillance. Using exome sequencing, we identified a *TP53* p.C242F mutation with CN-LOH in the AML sample (Figure 6A), then used deep, error-corrected sequencing to quantify mutation allele burden in bulk DNA across serial samples obtained prior to transformation. The *TP53* p.C242F mutation was first detectable at low abundance (VAF=0.17%) 4.5 years prior to transformation (Figure 6C). In addition to this pre-leukemic mutation, we also identified 17 additional mutations from 4 samples across 6.6 years of surveillance (ages 16-22), including *TP53* (n=7), *EIF6* (n=7), *CSNK1A1* (n=2), *PRPF8* (n=1), and *SBDS* (n=1)(Figure 6D). Throughout surveillance, the pre-leukemic *TP53* p.C242F clone was indistinguishable from other *TP53* mutated clones based on low VAF and relative stability across serial samples.

Bulk sequencing cannot reliably identify interval acquisition of *TP53* allelic imbalance in small clones. We therefore sequenced 20,214 single cells across 3 samples obtained 6.5, 4.5, and 0.5 years before clinical transformation in order to identify the earliest evidence of *TP53* CN-LOH. All *TP53* mutations detected by bulk sequencing were also observed using single cell DNA sequencing, but only the *TP53* p.C242F clone displayed evidence of clonal evolution with *TP53* LOH. Concordant with bulk sequencing data, the *TP53* p.C242F was first detectable 4.5 years prior to development of AML. The *TP53* p.C242F clone was initially present at low abundance (0.1%), with a balanced proportion of the heterozygous founding clone and the homozygous (CN-LOH) progression subclone. Subsequently, the CN-LOH subclone expanded selectively over the following 4 years prior to subsequent transformation. Other stable *TP53* mutations, including the most abundant p.R248W and p.L257R clones, defined independent clones and remained in the monoallelic state across 6.5 years of surveillance. These data indicate that development of *TP53* LOH events can precede frank transformation by several years and that single cell DNA sequencing enables detection of small clones defined by *TP53* LOH events.

## Discussion

We found that germline SBDS deficiency establishes a global fitness constraint that drives selection of somatic clones via two pathways with distinct mechanisms and different clinical consequences. A compensatory pathway with limited leukemic potential, mediated predominantly by EIF6 inactivation, enhances clone fitness by ameliorating the SDS ribosome defect. A maladaptive pathway with enhanced leukemic potential, driven by *TP53* inactivation, subverts normal tumor suppressor checkpoints without correcting the ribosome defect (Figure 7).

**Figure 7.**
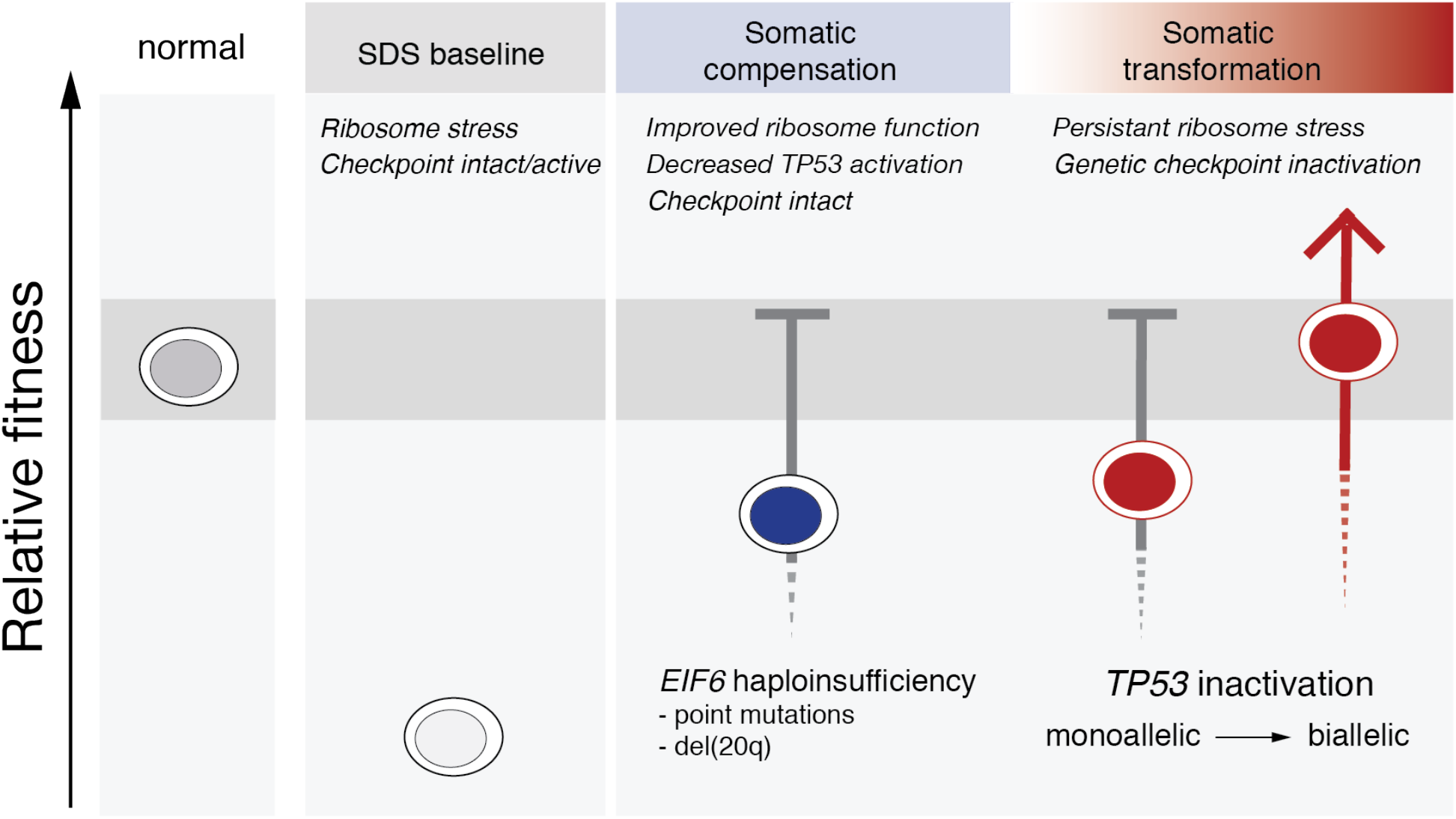
*TP53* and *EIF6* mutations define distinct pathways of somatic clonal progression and distinguish leukemia predisposition in SDS. Germline context drives separate compensatory and maladaptive somatic pathways of clonal evolution in patients with SDS. Germline *SBDS* mutations result in ribosomal stress which activate TP53 checkpoint pathways and promote bone marrow failure. *EIF6* mutations alleviate the underlying ribosome maturation defects which reduces p53 checkpoint activation and improves cell fitness. *TP53* mutations eliminate checkpoint pathways to improve relative fitness without improving the underlying ribosomal abnormalities, and promote the development of myeloid malignancies.

Through analysis of serial samples collected in children and young adults with SDS, we demonstrate that somatic clones are infrequent in the first several years of life, but approach ubiquity in the second decade. Many patients had multiple somatic mutations, but this high somatic mutation burden typically reflects a composite of multiple, genetically distinct and independently arising clones rather than a single clone with complex subclonal evolution. Most of these somatic clones in SDS patients carried a mutation in one of only four genes (*EIF6*, *TP53*, *PRPF8*, and *CSNK1A1*), and rarely involved genes commonly mutated in age-related CH (*DNMT3A*, *TET2*, *ASXL1*). These results provide genetic evidence that germline SBDS deficiency causes a global, disease-specific HSC fitness constraint that drives parallel development of somatic CH at an early age.

*EIF6* mutations have not previously been reported in human disease. We show that somatic *EIF6* mutations are common in SDS and that they cause functional compensation of SBDS deficiency by rescuing the SDS ribosome joining defect, improving translation, and reducing p53 activation. Using structural modeling and functional studies, we demonstrate that *EIF6* missense mutations exert these effects by either disrupting the binding interaction between EIF6 and the 60S subunit or by destabilizing the EIF6 protein. Our findings in SDS patients are consistent with prior studies in yeast, where mutations in the *EIF6* homolog *TIF6* were shown to attenuate the slow growth phenotype seen with deletion of the *SBDS* homolog *SDO1^27^*. Notably, several residues that we found to be recurrently mutated in SDS patients, including p.G14, p.G105, and p.N106, are invariant between human and Archaea and were found in a yeast genetic screen to cause reduced affinity for 60S subunits.

Normalization of germline-encoded hematopoietic defects through somatic reversion has been observed in inherited bone marrow failure syndromes^31–33^. In SDS, we found that somatic mutations mitigate the cellular consequences of SBDS deficiency via compensatory mechanisms without direct reversion or correction of the causative genetic lesion. Specifically, highly recurrent functional inactivation of EIF6 via multiple genetic mechanisms (point mutations and interstitial del(20q)) indicates that normalizing the functional ratio of SBDS:EIF6 protein in SDS hematopoietic cells enhances competitive fitness by improving ribosome maturation and translational capacity. *EIF6* alterations are not associated with leukemic transformation or *TP53* co-mutation within the same cell and were not found in patients with severe bone marrow failure, suggesting that functional correction of germline-encoded cellular defects may drive enhanced fitness of somatic clones without altering normal pathways of differentiation or oncogene protection. Our results support EIF6 as a potential therapeutic target in SDS patients, since pharmacologic inactivation of EIF6 could mimic genetic inactivation of *EIF6*, thereby reducing leukemia risk and improving hematopoietic function in SDS patients.

The presence, number, persistence, and allele abundance of somatic *TP53* mutations were not predictive of imminent leukemia risk in SDS patients with CH. Instead, our results indicate that progression of *TP53* mutated clones is driven by development of biallelic alterations of the *TP53* locus via deletion, CN-LOH, or point mutation. Importantly, we found that SDS patients can develop multiple, independent TP53 mutated clones and that serial monitoring by bulk sequencing fails to distinguish clinically significant subclonal changes in *TP53* allelic state. These findings support the integration of single cell DNA sequencing into surveillance strategies to identify patients with high risk clones.

In conclusion, our study defines a conceptual framework and strategy for rational surveillance in SDS patients. An improved ability to identify patients with high risk of developing leukemia has the potential to improve clinical outcomes by enabling preemptive intervention with curative therapies, such as allogeneic transplantation.

## Supporting information

Rosetta scripts

## Acknowledgements

This work is dedicated to the patients and families participating in the North American SDS Registry. We are grateful for assistance from the Dana Farber Cancer Institute Center for Cancer Genome Discovery (Aaron Thorner, Anwesha Nag), the Flow Cytometry Core facilities at Boston Children’s Hospital (Ronald Mathieu) and the Dana Farber Cancer Institute (Suzan Lazo), Dr. Soheil Meshinchi at the Fred Hutchinson Cancer Research Center, and the Boston Children’s Hospital Translab (Myriam Armant). Peripheral blood mobilized, cryopreserved CD34^+^ HSPCs from anonymous healthy donors were obtained from Fred Hutchinson Cancer Research Center, Seattle, Washington (NIDDK Cooperative Centers of Excellence in Hematology). This work was supported by the National Institutes of Health [R24 DK099808 (A.S., M.D.F.), 1RC2DK122533 (A.S. R.C.L., M.D.F.), K08CA204734 (R.C.L.)], the Butterfly Guild (R.C.L. and A.S.), the Edward P. Evans Foundation (A.L.K.) and the St. Baldrick’s Foundation (A.L.K.). Portions of this research were conducted on the O2 High Performance Computer Cluster, supported by the Research Computing Group, at Harvard Medical School. See http://rc.hms.harvard.edu for more information.

## Author Contributions

A.L.K, R.C.L and A.S conceived of the study, performed data acquisition, functional analysis, data interpretation and drafted the manuscript. C.B and J.B. performed in silico modeling of protein interactions. K.B., G.M.M., C.E.H. performed data acquisition and functional analysis. C.B. and S.B. performed data acquisition. K.C.M, E.F., A.G., M.M., D.D., J.M.G., T.N., A.A.B., A.V., J.M.L., J.C., I.H., J.N.H., E.S., N.H., J.C., B.W.L., A.L., W.S., P.C., M.N., J.R.E., K.W., R.H.H., S.K., M.D.F., S.M.D., contributed clinical phenotyping, data acquisition and clinical analysis of patient cohort. S.L., and E.A.W performed logistic and poisson regression modeling. C.J.G., N.C., R.H.K., S.L.C., and R.C.L. performed genomic analysis. All authors have given final approval of the submitted version and agree to be accountable for all aspects of the work related to the accuracy or integrity of the work.

## Declaration of Financial or Non-financial Competing Interests

J.R.E is a part of the speakers' Bureau: Jazz Pharmaceuticals; Astellas, Consultant: Jazz Pharmaceuticals. N.H. is a consultant for Novartis and Incyte. S.D. is a consultant for Alexion and Novartis. R.C.L. has received research funding from Jazz Pharmaceuticals, and consulting fees from Takeda Pharmaceuticals and bluebird bio.

## Online Materials and Methods

### Patients and Samples

Subjects provided written informed consent for protocols approved by the institutional review boards of Boston Children’s Hospital and Cincinnati Children’s Hospital, in accordance with the Declaration of Helsinki’s Ethical Principles of Medical Research Involving Human Subjects. All subjects provided informed consent prior to their participation in the study. Clinical criteria for SDS diagnosis were as previously described^34^.

### DNA Extraction

DNA was extracted from patient samples and patient fibroblasts using the QIAamp DNA Blood Mini kit (Qiagen, 51104) according to manufacturer instructions.

### Cell culture

Human leukemia cell lines (K562, isogenic K562 with a CRISPR-HDR corrected *TP53* allele constructed as previously described^35^), kindly provided by Benjamin Ebert (Dana Farber Cancer Institute) were maintained in RPMI media (Gibco, 11875-119) supplemented with 10% FBS, penicillin and streptomycin (Gibco). Primary cultures of bone marrow fibroblasts were established as previously described and maintained in Chang D media (Irvine Scientific, T105). Mobilized peripheral blood CD34+ and bone marrow mononuclear cells were maintained in GMP SCGM Serum-free Media (Cellgenix, 20802-0500) supplemented with 100ng/mL hSCF (Peprotech, 300-07), hTPO (Peprotech, 300-18), hFLT3-L (Peprotech, 300-19) and for bone marrow mononuclear cell culture, 20 ng/mL of IL-3 (R&D systems 203-IL-010/CF) was added. All cells lines were cultured at 37 °C under 5% CO2 and routinely screened for mycoplasma.^36^

### Colony Formation Assays

For methylcellulose colony formation assays, G-CSF mobilized peripheral blood CD34+ cells (Fred Hutch CCEH Core B) were resuspended in GMP SCGM Serum-free Media plus cytokines noted above (Cellgenix, 20802-0500) and allowed to recover for 36 hours. Then 90,000 BMMNC or 1,750 CD34+ cells were added to 3.5 ml of methylcellulose (Stem Cell Technologies, H4434). One ml was plated in triplicate wells of 6-well Smartdishes (Stem Cell Technologies, 27370). After 12 days of growth at 37°C/5% CO2, colonies were imaged, results blinded and then counted using STEMVision (Stem Cell Technologies). Counts were averaged for triplicate wells.

### Plasmids, Cloning and Site directed Mutagenesis

Gateway vectors containing EIF6 cDNA were obtained from the Harvard Plasmid Repository in closed format (clone ID HsCD00044644) or open format (clone ID HsCD00041550). Site directed mutagenesis was performed using the NEB Q5 site directed mutagenesis kit according to manufacturer instructions (New England Biolabs, E0554S) using primers listed in Supplementary Table 3. Gateway cloning was performed using LR clonase (Invitrogen, 11791-020) according to manufacturer instructions. Closed constructs were cloned into pRRL-SFFV-gwdest. Open constructs were cloned into constructs with V5 C-terminal tags: tetracycline inducible pLIX-403 or constitutive expression pLX304. pLIX-403 and pLX304^37^ were gifts from David Root (pLIX-403 is Addgene plasmid #41395, pLX304 is Addgene plasmid #25890). Tetracycline-inducible short hairpin RNAs targeting SBDS were made by annealing oligos and ligating with T4 ligase (New England Biolabs, M0202) into AgeI and EcoRI digested Tet-pLKO-puro. Tet-pLKO-puro was a gift from Dmitri Wiederschain (Addgene plasmid #21915)^38^.

### Immunofluorescence

Primary bone marrow fibroblasts from SDS patients were grown on coverslips in a six well plate at a density of 3 × 10^5^ cells/coverslip for 24 hours. Cells were washed with PBS (Gibco), fixed with 4% (w/v) paraformaldehyde (MilliporeSigma) for 10 minutes at room temperature, washed 3 times with PBS, permeabilized with 0.2% Triton X-100 (VWR) for 5 minutes at room temperature, washed 3 times with PBS, and blocked for 30 minutes at room temperature in solution with 3% BSA (MilliporeSigma) before 30 minute incubation with primary antibody at room temperature (Fibrillarin, Cell Signaling, clone C13C3), (V5, MBL, Clone OZA3). Coverslips were washed 3 times with 1% Triton(VWR) in PBS before incubation with secondary antibody (AlexaFluor 594 donkey anti–rabbit IgG (Fisher Scientific, A21207) or AlexaFluor 488 donkey anti-mouse, (Life Technologies, A-21202)). Cells were mounted with mounting medium containing DAPI (Vector Laboratories, H-1200) for nuclear counterstaining and were imaged with a Nikon Eclipse 90i microscope.

### Polysome Profiling

Ribosomal subunits were separated by sucrose density gradients as described^28^. Briefly, 4 × 10^6^ fibroblasts (80 % confluence) or 2.5 ×10^6^ K562 (1 million/mL) were treated with cycloheximide at final concentration of 100 μg/mL for 10 min at 37 °C before harvesting. Cells were then lysed in lysis buffer (20 mM HEPES pH 7.4, 100 mM KCl, 10 mM MgCl2, 1% [w/v] NP-40, 1% [w/v] deoxycholate, 100 μg/mL cycloheximide,1mM DTT, 200 μg/mL heparin, with complete EDTA-free protease inhibitors (Roche), 1.4uM pepstatin A, and 40 U/mL Rnasin (Promega N2115) and incubated for 10 min on ice. Lysates were cleared in a microcentrifuge at 4 °C. Equal amounts were applied to a 5-50 % (w/v) sucrose gradient in gradient buffer (10 mM HEPES pH 7.4, 50 mM KCl, 5 mM MgCl2, 100 μg/mL cycloheximide) and centrifuged (Beckman SW55.1 rotor at 44,000 rpm for 1 hour and 15 minutes at 4 °C). The sucrose gradient was made using a Biocomp Gradient Master. Centrifugation samples were unloaded using a Brandel gradient fractionator, polysome profiles detected at 254nM absorbance and area under the curve for 80S and 60S peaks quantitated using Peakchart version 2.08 (Brandel). 0.1 mL fractions collected into Laemmli sample buffer and separated on SDS-PAGE gels and transferred to PVDF membranes for immunoblotting.

### qRT–PCR

RNA was isolated following manufacturer’s instructions for RNeasy Plus Mini Kit (QIAGEN Inc., 44134). RNA was eluted in 30 μl of water. We used 200 ng to 1 μg of RNA for reverse transcription with Superscript III First Strand Synthesis using oligo-dT primer (Invitrogen, 18080051). For qPCR analysis, cDNAs were diluted threefold in MilliQ water. Quantitative PCR was performed with iTaq Universal SYBR Green Supermix (Bio-Rad, 1725125). The linear range of amplification for each primer pair was confirmed by serial dilution of genomic DNA from K562 cells. Reactions were carried out in triplicate in a 7500 Fast Real-Time PCR System (Applied Biosystems) and analyzed using the ΔΔCT method^39^. The primer sequences are shown in Supplementary Table 3.

### Virus Production and Titration

Transfection of 293T cells was performed as described^40^. Lentiviral vector supernatants were generated by cotransfecting lentiviral transfer vectors (pRRL-SFFV-gwdest, pLIX-403 (Addgene plasmid #41395), pLX304 (Addgene plasmid #25890), tet-pLKO-puro (Addgene plasmid #21915), SMARTvector-human-shEIF6 (Dharmacon V3SH11243-07EG3692), SMARTvector-hCMV-shTP53-TurboGFP (Dharmacon V3SH11243-00EG7157), SMARTvector-hCMV-shTP53-TurboRFP (Dharmacon V3SH11243-07EG7157), pL40C.SFFV.eGFP.miR30N.PRE-shLUC^41^ (CGCTGAGTACTTCGAAATGTC) or pL40C.SFFV.eGFP.miR30N.PRE-shSBDS with packaging plasmids psPAX2 (Addgene plasmid #12260) and pMD2.G (Addgene plasmid #12259) using PEI reagent (Polysciences #23966-2). Supernatants were collected, filtered through a 0.45 μm membrane (ThermoFisher, 165-0045), and subsequently concentrated by ultracentrifugation at 10,000 rpm for 10 hours in a Beckmann XL-90 centrifuge using SW-28 swinging buckets. To determine the titer, HT1080 cells were infected with the virus in the presence of 8 μg/ml polybrene (Santa Cruz, SC134220) and analyzed 48 hours post-transduction by fluorescence-activated cell sorting for GFP expression.

### Single-cell DNA sequencing

We designed a custom panel covering 7 genes implicated in SDS or sporadic clonal hematopoiesis and 43 single nucleotide polymorphism loci on chromosomes 7, 17, and 20 (Supplementary Table 4). Libraries were generated from cyropreserved or fresh bone marrow mononuclear cells with the Mission Bio Tapestri Single-cell DNA custom Kit according to manufacturer’s instruction (Mission Bio) with the following modifications: concentration of cell input was increased by 15% and library PCR cycles were increased by one cycle. Libraries were pooled in equimolar concentration and sequenced on a NovaSeq (Illumina) on a 150 base pair paired end run. Analysis including data filtering and visualization was conducted using Tapestri Insights 2.2 (Mission Bio).

### Whole Exome Sequencing

Prior to library preparation, DNA was fragmented (Covaris sonication) to 250 bp and further purified using Agentcourt AMPure XP beads. Size-selected DNA was then ligated to specific adapters during automated library preparation (SPRIworks, Beckman-Coulter). Libraries were pooled and sequenced on an Illumina MiSeq to estimate the concentration based on the number of barcode reads per sample). Library construction is considered successful if the yield is ≥ 250 ng. Libraries were pooled in equal mass to a total of 750 ng for SureSelect Human All Exon V5 enrichment using the Agilent SureSelect hybrid capture kit. Captures were further pooled and sequenced on HiSeq2500 or HiSeq3000 (Illumina). Pooled sample reads were de-convoluted (de-multiplexed) and sorted using the Picard tools^42^. Reads were aligned to the reference sequence b37 edition from the Human Genome Reference Consortium using “bwa aln” (http://bio-bwa.sourceforge.net/bwa.shtml) using the following parameters “-q 5 -l 32 -k 2 -o 1” and duplicate reads were identified and removed using the Picard tools. The alignments were further refined using the GATK tool for localized realignment around indel sites. Recalibration of the quality scores was also performed using GATK tools^43,44^. Metrics for the representation of each sample were generated on the unaligned reads after sorting on the barcode. Fingerprinting analysis was performed using 44 polymorphic loci to identify if the aggregation pairing strategy was performed appropriately. Picard Tools GenotypeConcordance was used to calculate the concordance that a given test sample matches the sample being considered. This was performed on all pairwise combinations of samples in the cohort. The output of the pairwise comparisons was then mapped to a concordance matrix, where concordance values above 4 standard deviations of the median concordance value for the cohort indicated a high likelihood that the samples match. Samples can match for reasons other than being from the same individual, so potential matches are manually reviewed where applicable.

For bone marrow samples, median of mean target coverage = 136x, range 101-198x. Mutation analysis for single nucleotide variants (SNV) was performed using MuTect v1.1.4 and annotated by Variant Effect Predictor (VEP). We used the SomaticIndelDetector tool that is part of the GATK for indel calling. Variant Effect Predictor v79 is used for annotating the variants. MuTect was run in paired mode with bone marrow aspirate and cultured fibroblast samples from each subject. Variants that affected protein coding regions underwent further filtering/classification based on frequency in the gnomAD, ESP, and COSMIC (version 80) databases. Variants that affect protein coding regions were flagged as “REVIEW_REQUIRED”, if the frequency of the variant is less than or equal to 1% in all gnomAD and ESP populations or if the frequency of the variant is greater than 1% and less than or equal to 10% in all gnomAD and ESP populations and present in “COSMIC” database at least 2 times. Variants were flagged as “NO_REVIEW_GERMLINE_FILTER” if the frequency of the variant is between 1% and less than or equal to 10% in all gnomAD and ESP populations and not present in “COSMIC” database at least 2 times or if the frequency of the variant is greater than 10% in any gnomAD and ESP populations. Variants with frequency greater than 10% in any gnomAD or ESP population were considered to be a common SNP irrespective of presence in the COSMIC database.

### Copy number analysis

To obtain raw copy-number estimates across the genome of each sample, the number of unique templates mapping to each exome target region (padded by 250 bp) was extracted from the BAM file. The raw estimates were normalized against coverage obtained from a panel of diploid normal samples. A subset of targets was removed based on estimates of mean total copy-ratio and the standard deviation of copy-ratio estimates within a panel of diploid normal samples. The resulting total copy-ratio profiles were then segmented using an adaptation of the circular binary segmentation algorithm, which includes information from all patient samples when segmenting. Subsequently, the allele-specific copy number was estimated by examining the template counts supporting alternative and reference alleles at germline heterozygous SNP sites within the 1000 Genomes Phase 3 variants. Of the 1000 Genomes Phase 3 variants, a patient was considered heterozygous at a given locus based on the number of reference and alternative template counts observed in all patient samples. The allele-specific template counts were then used to infer allele-specific copy ratios as described previously serving as input into ABSOLUTE v.1.4, which jointly estimated the fraction of cancer cells, cancer ploidy and absolute allelic copy numbers across the genome.

### Targeted Deep Sequencing

We selected 55 genes for targeted sequencing based on their recurrent alteration in SDS exome cohort and myeloid malignancies^13^ (Supplementary Table 1). We included 48 single nucleotide polymorphisms (SNPs) for establishing subject concordance of serial samples. *Library Construction:* An aliquot of genomic DNA (50-200ng in 50μL) was used as the input into DNA fragmentation (aka shearing). Shearing was performed acoustically using a Covaris focused-ultrasonicator, targeting 150bp fragments. Library preparation was performed using a commercially available kit provided by KAPA Biosystems (KAPA HyperPrep Kit with Library Amplification product KK8504) and IDT’s duplex UMI adapters. The libraries were then paired with unique 8-base dual index sequences embedded within the p5 and p7 primers (purchased from IDT) added during PCR. Enzymatic clean-ups are performed using Beckman Coulter AMPure XP beads with elution volumes reduced to 30μL to maximize library concentration. In addition, during the post-enrichment SPRI cleanup, elution volume was reduced to 30μL to maximize library concentration, and a vortexing step was added to maximize the amount of template eluted. *Post Library Construction Quantification and Normalization:* Library quantification was performed using the Invitrogen Quant-It broad range dsDNA quantification assay kit (Thermo Scientific Catalog: Q33130) with a 1:200 PicoGreen dilution. Following quantification, each library is normalized to a concentration of 35 ng/μL, using Tris-HCl, 10mM, pH 8.0. *In-solution hybrid selection:* After library construction, hybridization and capture are performed using the relevant components of IDT’s XGen hybridization and wash kit and following the manufacturer’s suggested protocol, with several exceptions. A set of 12-plex pre-hybridization pools are created. These pre-hybridization pools are created by equivolume pooling of the normalized libraries, Human Cot-1 and IDT XGen blocking oligos. The pre-hybridization pools undergo lyophilization using the Biotage SPE-DRY. Post lyophilization, custom exome bait (TWIST Biosciences) along with hybridization mastermix is added to the lyophilized pool prior to resuspension. Samples are incubated overnight. Library normalization and hybridization setup are performed on a Hamilton Starlet liquid handling platform, while target capture is performed on the Agilent Bravo automated platform. Post capture, a PCR is performed to amplify the capture material. After post-capture enrichment, library pools are quantified using qPCR (automated assay on the Agilent Bravo), using a kit purchased from KAPA Biosystems with probes specific to the ends of the adapters. Based on qPCR quantification, pools are normalized using a Hamilton Starlet to 2nM and sequenced using Illumina sequencing technology. *Cluster amplification and sequencing:* Cluster amplification of library pools was performed according to the manufacturer’s protocol (Illumina) using Exclusion Amplification cluster chemistry and HiSeq X flowcells. Flowcells were sequenced on v2 Sequencing-by-Synthesis chemistry for HiSeq X flowcells. The flowcells are then analyzed using RTA v.2.7.3 or later. Each pool of whole genome libraries was run on paired 151bp runs, reading the dual-indexed sequences to identify molecular indices and sequenced across the number of lanes needed to meet coverage for all libraries in the pool.

### Variant calling pipeline

Reads are aligned with bwa-mem 0.7.15. Duplex consensus reads are called with fgbio 1.0 and realigned using bwa-mem. Consensus reads are required to have reads from both families αβ and βα, and consensus reads with Ns in excess of 5% of bases are discarded. Read one and two are soft-clipped from the 5' end by 10 bases to reduce errors due to end repair. Single nucleotide and small insertion and deletion calling was performed with samtools-0.1.18 mpileup and Varscan 2.2.3. Variants were annotated to include information about cDNA and amino acid changes, sequence depth, number and percentage of reads supporting the variant allele, population allele frequency in 1000 Genomes release 2.2.2^45^, the Genome Aggregation Database (gnomAD)^46^, and presence in Catalogue of Somatic Mutations in Cancer (COSMIC), version 64.6^47^. Variants were excluded if they had fewer than 3 total duplex reassembled alternate reads at the position or had variant allele fraction < 0.1%, fell outside of the target coordinates, had excessive read strand bias, had excessive number of calls in the local region, caused synonymous changes, or were recurrent small insertions/deletions at low variant allele fraction adjacent to homopolymer repeat regions. Somatic status was determined using cultured fibroblast DNA as a germline reference tissue comparator. Individual single nucleotide substitutions and small insertions or deletions were evaluated as candidate drivers of MDS or bone marrow failure based on gene-specific characteristics, then curated manually and classified as MDS driver mutations or pathogenic bone marrow failure mutations based on genetic criteria and literature review^13,48,49^. Variant level details are available in Supplementary Table 5. All interpretation of variants was blinded to clinical characteristics and thus agnostic to variables including age, sex, diagnosis, treatment status, and clinical outcomes; the genetic analysis was completed and locked prior to merging with any clinical data.

### Growth Competition Assay

K562 *TP53* corrected cells containing puromycin selectable doxycycline inducible shRNA targeting SBDS (target sequence GCTTGGATGATGTTCCTGATT) were transduced with lentivirus encoding constitutively expressed EIF6-RFP mutant or wild-type constructs as noted. After 48 hours, cells were admixed at a ratio of 1:1 and subjected to flow cytometry at indicated times on a Fortessa HTS flow cytometer (BD Biosciences). Cells maintained in puromycin (Mirus, 2ug/mL) throughout the experiment. Analysis was performed with FacsDIVA software (BD Biosciences).

### Immunoblotting

Cells were lysed in RIPA buffer (MilliporeSigma) supplemented with protease inhibitors (Inhibitab, Roche). Protein concentrations were determined by colorimetric assay (BCA Protein, Thermo Fisher Scientific), and 20-40 μg of protein was loaded on 12% SDS-PAGE gels and blotted on a PVDF membrane (MilliporeSigma, IPVH00010). The membranes were blocked with 5% nonfat dry milk (VWR) diluted in Tris-buffered saline (Teknova, T1680) with 1% Tween-20 (VWR, M147-1L). Primary antibodies SBDS^50^, GAPDH (Cell Signaling Technology, clone 14C10), eIF6 (Cell Signaling, clone D16E9), p53 Calbiochem, OP43), RPL3 (Abcam, ab241412), Ubiquitin (Cell signaling, 3933), V5 tag (Abcam, ab15828), Vinculin (Invitrogen, clone 42H89L44), were incubated overnight at 4°C. After washing with TBS-T, membranes were incubated with HRP-conjugated secondary antibodies ECL anti-rabbit IgG (GE Healthcare, NA934V) and ECL anti-mouse (GE Healthcare, NA931V) and developed using SuperSignal West Pico Chemiluminescent substrate (Thermo Fisher Scientific, 34094). Detection of bands was conducted in the Amersham Imager 600 (GE Healthcare).

### OP-Puro Incorporation

OP-Puro (Medchem Source; Life Technologies, C10459; 50 μM final concentration) was added to the culture medium for 3 hours and incubated in a 37° incubator. Cells were removed from wells and washed twice in Ca2+ and Mg2+ free phosphate buffered saline (PBS) + cycloheximide. Cells were fixed in 0.5ml of 1% paraformaldehyde in PBS for 10 minutes, then permeabilized in PBS supplemented with 3% fetal bovine serum and 0.1% saponin for 5 minutes at room temperature. The azide-alkyne cycloaddition was performed using the Click-iT Cell Reaction Buffer Kit (Life Technologies, C10458) and azide conjugated to Alexa Fluor 647 (Life Technologies, C10458) at 5μM final concentration for 30 minutes. The cells were washed twice in PBS and passed through a filter top tube prior to being analyzed by flow cytometry.

### Statistical analysis

Graphpad Prism version 8 and SAS 9.4 (SAS Institute, Cary NC) was used to analyze results and create graphs. Comparison of polysome 80S/60S area under the curve and CFUs represent 2-tailed student t-test. Fisher’s exact test is used to assess the association between presence of mutations and patient characteristics. Wilcoxon rank sum test is used to assess the association between number of mutations and patient characteristics. Results are considered significantly associated with outcome if p-values <0.05 and marginally associated with outcome if p<0.10.

### EIF6 Model and Mutational Analysis

A human EIF6 structural model was generated using Robetta^51,52^, and the structure was then further refined in Rosetta using the FastRelax algorithm^53^ with the *Rosetta-ICO* energy function^54^. Individual point mutations were evaluated for the predicted change in protein stability (ΔΔ*G*_mutation_) by introducing point mutations into the EIF6 structural model and calculating the change in energy (i.e. Rosetta total_score) relative to the native structure. The residues at the interface between EIF6 and RPL23 are 100% conserved between our model of human EIF6 and *D. discoideum* EIF6; thus, PDB ID 5ANB^22^ was refined in the same way and used for the ΔΔ*G*_bind_ calculations, which were performed using the Flex ddG method^23^. All scripts that were used for refinement and analysis are provided in the supplementary material. Conservation scores were calculated using the ConSurf server ^55^, and structural images were generated using PyMOL.

