## Supplementary Figures for "Distinct genetic pathways define pre-leukemic and compensatory clonal hematopoiesis in Shwachman-Diamond syndrome"

**Supplementary Data**

**Supplementary Figure 1**

A

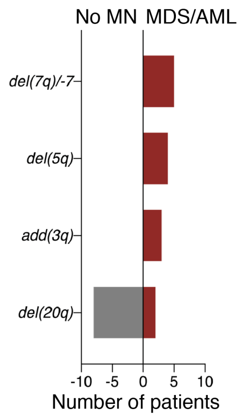


**Supplementary Figure 1: Recurrent structural alterations found in patient cohort with or without myeloid neoplasm.** Shown are the indicated somatic copy number alterations in the exome cohort without (grey bar) or with (red bars) myeloid neoplasm.

**Supplementary Figure 2**

A

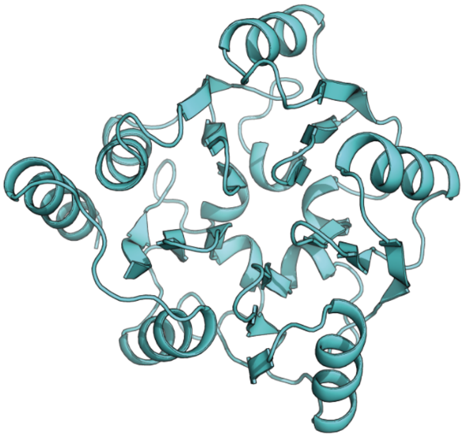

**Supplementary Figure 2: Homology model of human EIF6 protein.** Propeller like structural model of human EIF6 contains the five conserved ββαβ motifs.

**Supplementary Figure 3**

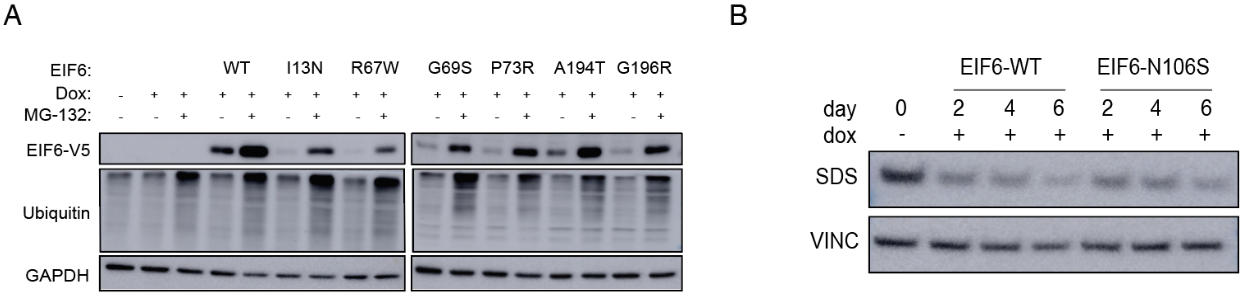


**Supplementary Figure 3: EIF6 protein degradation is proteasome dependent**. **A,** Immunoblot of K562 cells expressing doxycycline-inducible V5-tagged wild type EIF6 (WT) or mutant EIF6 cDNAs and treated with MG-132 proteasome inhibitor as indicated. The blots were probed with antibodies against the V5 tag, ubiquitin, or GAPDH. Ubiquitin serves as control for MG-132 activity and GAPDH as loading control. **B**, Western blot of K562 cells expressing inducible shRNAs targeting SBDS at timepoints indicated.

**Supplementary Figure 4**

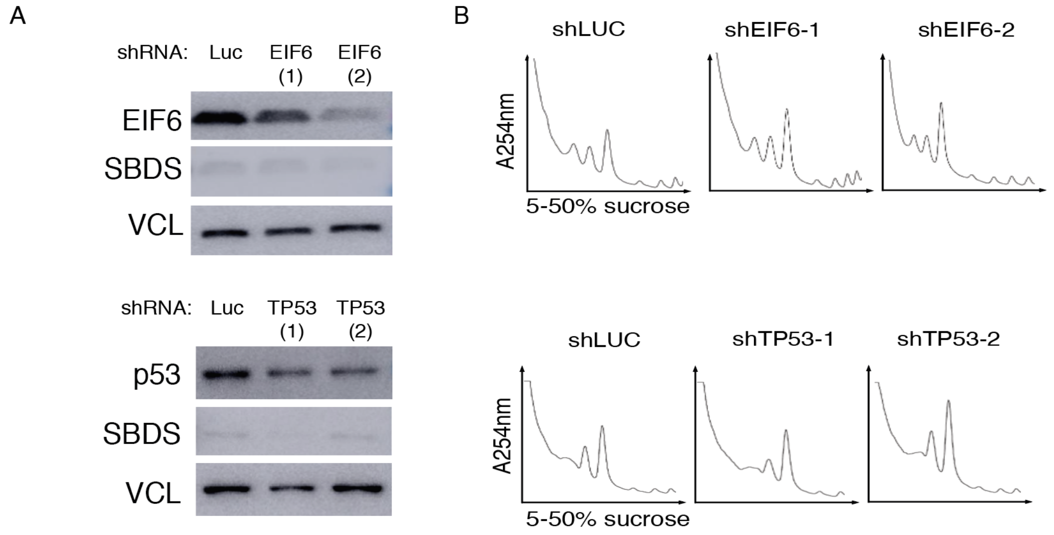


**Supplementary Figure 4: Polysome profiles of *EIF6* or *TP53* deficient patient-derived SDS fibroblasts**. **A,** EIF6, SBDS and vinclulin immunoblots of primary SDS patient-derived fibroblasts transduced with shRNAs targeting luciferase control, EIF6 (top panel) or TP53 (bottom panel). **B,** Polysome profiles of cells in panel A.
