## Supplementary material for "Distinct genetic pathways define pre-leukemic and compensatory clonal hematopoiesis in Shwachman-Diamond syndrome": Rosetta scripts: run.sbatch

```
#!/bin/bash
#SBATCH -c 1                               # Request one core
#SBATCH -N 1                               # Request one node (if you request more than one core with -c, also using
#SBATCH -t 0-00:15                         # Runtime in D-HH:MM format
#SBATCH -p short                           # Partition to run in
#SBATCH --mem=1G                           # Memory total in MB (for all cores)
#SBATCH --array=0-99
#SBATCH -o runlogs/%j.out
#SBATCH -e runlogs/%j.err
#SBATCH --mail-type=END                    # Type of email notification- BEGIN,END,FAIL,ALL
#SBATCH --mail-user=

DIRECTORY=/home/jwb29/eif6/prep_structure
PDB=${DIRECTORY}/full_model_18244_1.pdb
SCRIPT=${DIRECTORY}/fastrelax.xml
OUTPUT=${DIRECTORY}/output/${SLURM_ARRAY_JOB_ID}_${SLURM_ARRAY_TASK_ID}.silent
SUFFIX=_${SLURM_ARRAY_JOB_ID}_${SLURM_ARRAY_TASK_ID}

/n/data1/bch/hemonc/bahl/samaneh/Rosetta/main/source/bin/rosetta_scripts.default.linuxgccrelease \
	-beta \
	-linmem_ig 10 \
	-in:file:s ${PDB} \
	-parser:protocol ${SCRIPT} \
	-out:file:silent ${OUTPUT} \
	-out:suffix ${SUFFIX} \
	-nstruct 1 \
```
