## Supplementary material for "Distinct genetic pathways define pre-leukemic and compensatory clonal hematopoiesis in Shwachman-Diamond syndrome": Rosetta scripts: analyze_flex_ddG.py

```
#!/usr/bin/python3

import sys
import os
import sqlite3
import shutil
import tempfile
from pprint import pprint
import pandas as pd
import numpy as np
import re
import datetime
import sys
import collections
import threading

rosetta_output_file_name = 'rosetta.out'
output_database_name = 'ddG.db3'
trajectory_stride = 5
script_output_folder = 'analysis_output'

zemu_gam_params = {
    'fa_sol' :      (6.940, -6.722),
    'hbond_sc' :    (1.902, -1.999),
    'hbond_bb_sc' : (0.063,  0.452),
    'fa_rep' :      (1.659, -0.836),
    'fa_elec' :     (0.697, -0.122),
    'hbond_lr_bb' : (2.738, -1.179),
    'fa_atr' :      (2.313, -1.649),
}

def gam_function(x, score_term = None ):
    return -1.0 * np.exp( zemu_gam_params[score_term][0] ) + 2.0 * np.exp( zemu_gam_params[score_term][0] ) / ( 1.0 + np.exp( -1.0 * x * np.exp( zemu_gam_params[score_term][1] ) ) )

def apply_zemu_gam(scores):
    new_columns = list(scores.columns)
    new_columns.remove('total_score')
    scores = scores.copy()[ new_columns ]
    for score_term in zemu_gam_params:
        assert( score_term in scores.columns )
        scores[score_term] = scores[score_term].apply( gam_function, score_term = score_term )
    scores[ 'total_score' ] = scores[ list(zemu_gam_params.keys()) ].sum( axis = 1 )
    scores[ 'score_function_name' ] = scores[ 'score_function_name' ] + '-gam'
    return scores

def rosetta_output_succeeded( potential_struct_dir ):
    path_to_rosetta_output = os.path.join( potential_struct_dir, rosetta_output_file_name )
    if not os.path.isfile(path_to_rosetta_output):
        return False

    db3_file = os.path.join( potential_struct_dir, output_database_name )
    if not os.path.isfile( db3_file ):
        return False

    success_line_found = False
    no_more_batches_line_found = False
    with open( path_to_rosetta_output, 'r' ) as f:
        for line in f:
            if line.startswith( 'protocols.jd2.JobDistributor' ) and 'reported success in' in line:
                success_line_found = True
            if line.startswith( 'protocols.jd2.JobDistributor' ) and 'no more batches to process' in line:
                no_more_batches_line_found = True

    return no_more_batches_line_found and success_line_found

def find_finished_jobs( output_folder ):
    return_dict = {}
    job_dirs = [ os.path.abspath(os.path.join(output_folder, d)) for d in os.listdir(output_folder) if os.path.isdir( os.path.join(output_folder, d) )]
    for job_dir in job_dirs:
        completed_struct_dirs = []
        for potential_struct_dir in sorted([ os.path.abspath(os.path.join(job_dir, d)) for d in os.listdir(job_dir) if os.path.isdir( os.path.join(job_dir, d) )]):
            if rosetta_output_succeeded( potential_struct_dir ):
                completed_struct_dirs.append( potential_struct_dir )
        return_dict[job_dir] = completed_struct_dirs

    return return_dict

def get_scores_from_db3_file(db3_file, struct_number, case_name):
    conn = sqlite3.connect(db3_file)
    conn.row_factory = sqlite3.Row
    c = conn.cursor()

    num_batches = c.execute('SELECT max(batch_id) from batches').fetchone()[0]

    scores = pd.read_sql_query('''
    SELECT batches.name, structure_scores.struct_id, score_types.score_type_name, structure_scores.score_value, score_function_method_options.score_function_name from structure_scores
    INNER JOIN batches ON batches.batch_id=structure_scores.batch_id
    INNER JOIN score_function_method_options ON score_function_method_options.batch_id=batches.batch_id
    INNER JOIN score_types ON score_types.batch_id=structure_scores.batch_id AND score_types.score_type_id=structure_scores.score_type_id
    ''', conn)

    def renumber_struct_id( struct_id ):
        return trajectory_stride * ( 1 + (int(struct_id-1) // num_batches) )

    scores['struct_id'] = scores['struct_id'].apply( renumber_struct_id )
    scores['name'] = scores['name'].apply( lambda x: x[:-9] if x.endswith('_dbreport') else x )
    scores = scores.pivot_table( index = ['name', 'struct_id', 'score_function_name'], columns = 'score_type_name', values = 'score_value' ).reset_index()
    scores.rename( columns = {
        'name' : 'state',
        'struct_id' : 'backrub_steps',
    }, inplace=True)
    scores['struct_num'] = struct_number
    scores['case_name'] = case_name

    conn.close()

    return scores

def process_finished_struct( output_path, case_name ):
    db3_file = os.path.join( output_path, output_database_name )
    assert( os.path.isfile( db3_file ) )
    struct_number = int( os.path.basename(output_path) )
    scores_df = get_scores_from_db3_file( db3_file, struct_number, case_name )

    return scores_df

def calc_ddg( scores ):
    total_structs = np.max( scores['struct_num'] )

    nstructs_to_analyze = set([total_structs])
    for x in range(10, total_structs):
        if x % 10 == 0:
            nstructs_to_analyze.add(x)
    nstructs_to_analyze = sorted(nstructs_to_analyze)

    all_ddg_scores = []
    for nstructs in nstructs_to_analyze:
        ddg_scores = scores.loc[ ((scores['state'] == 'unbound_mut') | (scores['state'] == 'bound_wt')) & (scores['struct_num'] <= nstructs) ].copy()
        for column in ddg_scores.columns:
            if column not in ['state', 'case_name', 'backrub_steps', 'struct_num', 'score_function_name']:
                ddg_scores.loc[:,column] *= -1.0
        ddg_scores = ddg_scores.append( scores.loc[ ((scores['state'] == 'unbound_wt') | (scores['state'] == 'bound_mut')) & (scores['struct_num'] <= nstructs) ].copy() )
        ddg_scores = ddg_scores.groupby( ['case_name', 'backrub_steps', 'struct_num', 'score_function_name'] ).sum().reset_index()

        if nstructs == total_structs:
            struct_scores = ddg_scores.copy()

        ddg_scores = ddg_scores.groupby( ['case_name', 'backrub_steps', 'score_function_name'] ).mean().round(decimals=5).reset_index()
        new_columns = list(ddg_scores.columns.values)
        new_columns.remove( 'struct_num' )
        ddg_scores = ddg_scores[new_columns]
        ddg_scores[ 'scored_state' ] = 'ddG'
        ddg_scores[ 'nstruct' ] = nstructs
        all_ddg_scores.append(ddg_scores)

    return (pd.concat(all_ddg_scores), struct_scores)

def calc_dgs( scores ):
    l = []

    total_structs = np.max( scores['struct_num'] )

    nstructs_to_analyze = set([total_structs])
    for x in range(10, total_structs):
        if x % 10 == 0:
            nstructs_to_analyze.add(x)
    nstructs_to_analyze = sorted(nstructs_to_analyze)

    for state in ['mut', 'wt']:
        for nstructs in nstructs_to_analyze:
            dg_scores = scores.loc[ (scores['state'].str.endswith(state)) & (scores['state'].str.startswith('unbound')) & (scores['struct_num'] <= nstructs) ].copy()
            for column in dg_scores.columns:
                if column not in ['state', 'case_name', 'backrub_steps', 'struct_num', 'score_function_name']:
                    dg_scores.loc[:,column] *= -1.0
            dg_scores = dg_scores.append( scores.loc[ (scores['state'].str.endswith(state)) & (scores['state'].str.startswith('bound')) & (scores['struct_num'] <= nstructs) ].copy() )
            dg_scores = dg_scores.groupby( ['case_name', 'backrub_steps', 'struct_num', 'score_function_name'] ).sum().reset_index()
            dg_scores = dg_scores.groupby( ['case_name', 'backrub_steps', 'score_function_name'] ).mean().round(decimals=5).reset_index()
            new_columns = list(dg_scores.columns.values)
            new_columns.remove( 'struct_num' )
            dg_scores = dg_scores[new_columns]
            dg_scores[ 'scored_state' ] = state + '_dG'
            dg_scores[ 'nstruct' ] = nstructs
            l.append( dg_scores )
    return l

def analyze_output_folder( output_folder ):
    # Pass in an outer output folder. Subdirectories are considered different mutation cases, with subdirectories of different structures.
    finished_jobs = find_finished_jobs( output_folder )
    if len(finished_jobs) == 0:
        print( 'No finished jobs found' )
        return

    ddg_scores_dfs = []
    struct_scores_dfs = []
    for finished_job, finished_structs in finished_jobs.items():
        inner_scores_list = []
        for finished_struct in finished_structs:
            inner_scores = process_finished_struct( finished_struct, os.path.basename(finished_job) )
            inner_scores_list.append( inner_scores )
        scores = pd.concat( inner_scores_list )
        ddg_scores, struct_scores = calc_ddg( scores )
        struct_scores_dfs.append( struct_scores )
        ddg_scores_dfs.append( ddg_scores )
        ddg_scores_dfs.append( apply_zemu_gam(ddg_scores) )
        ddg_scores_dfs.extend( calc_dgs( scores ) )

    if not os.path.isdir(script_output_folder):
        os.makedirs(script_output_folder)
    basename = os.path.basename(output_folder)

    pd.concat( struct_scores_dfs ).to_csv( os.path.join(script_output_folder, basename + '-struct_scores_results.csv' ) )

    df = pd.concat( ddg_scores_dfs )
    df.to_csv( os.path.join(script_output_folder, basename + '-results.csv') )

    display_columns = ['backrub_steps', 'case_name', 'nstruct', 'score_function_name', 'scored_state', 'total_score']
    for score_type in ['mut_dG', 'wt_dG', 'ddG']:
        print( score_type )
        print( df.loc[ df['scored_state'] == score_type ][display_columns].head( n = 20 ) )
        print( '' )

if __name__ == '__main__':
    for folder_to_analyze in sys.argv[1:]:
        if os.path.isdir( folder_to_analyze ):
            analyze_output_folder( folder_to_analyze )
```
