## Supplementary material for "Distinct genetic pathways define pre-leukemic and compensatory clonal hematopoiesis in Shwachman-Diamond syndrome": Rosetta scripts: eif6.py

```
#!/usr/bin/python

from __future__ import print_function

import socket
import sys
import os
import subprocess

use_multiprocessing = True
if use_multiprocessing:
    import multiprocessing
    max_cpus = 4 # We might want to not run on the full number of cores, as Rosetta take about 2 Gb of memory per instance

###################################################################################################################################################################
# Important: The variables below are set to values that will make the run complete faster (as a tutorial example), but will not give scientifically valid results.
#            Please change them to the "normal" default values before a real run.
###################################################################################################################################################################

rosetta_scripts_path = os.path.expanduser("/n/data1/bch/hemonc/bahl/samaneh/Rosetta/main/source/bin/rosetta_scripts.default.linuxgccrelease")
nstruct = 35 # Normally 35
max_minimization_iter = 5000 # Normally 5000
abs_score_convergence_thresh = 1.0 # Normally 1.0
number_backrub_trials = 35000 # Normally 35000
backrub_trajectory_stride = 35000 # Can be whatever you want, if you would like to see results from earlier time points in the backrub trajectory. 7000 is a reasonable number, to give you three checkpoints for a 35000 step run, but you could also set it to 35000 for quickest run time (as the final minimization and packing steps will only need to be run one time).
path_to_script = 'ddG-backrub.xml'

def run_flex_ddg( name, input_path, input_pdb_path, chains_to_move, nstruct_i ):
    output_directory = os.path.join( 'output', os.path.join( name, '%02d' % nstruct_i ) )
    if not os.path.isdir(output_directory):
        os.makedirs(output_directory)

    flex_ddg_args = [
        os.path.abspath(rosetta_scripts_path),
        "-s %s" % os.path.abspath(input_pdb_path),
        '-parser:protocol', os.path.abspath(path_to_script),
        '-parser:script_vars',
        'chainstomove=' + chains_to_move,
        'mutate_resfile_relpath=' + os.path.abspath( os.path.join( input_path, 'nataa_mutations.resfile' ) ),
        'number_backrub_trials=%d' % number_backrub_trials,
        'max_minimization_iter=%d' % max_minimization_iter,
        'abs_score_convergence_thresh=%.1f' % abs_score_convergence_thresh,
        'backrub_trajectory_stride=%d' % backrub_trajectory_stride ,
        '-beta',
        '-in:file:fullatom',
        '-ignore_unrecognized_res',
        '-ignore_zero_occupancy false',
        '-ex1',
        '-ex2',
    ]

    log_path = os.path.join(output_directory, 'rosetta.out')

    print( 'Running Rosetta with args:' )
    print( ' '.join(flex_ddg_args) )
    print( 'Output logged to:', os.path.abspath(log_path) )
    print()

    outfile = open(log_path, 'w')
    process = subprocess.Popen(flex_ddg_args, stdout=outfile, stderr=subprocess.STDOUT, close_fds = True, cwd = output_directory)
    returncode = process.wait()
    outfile.close()

if __name__ == '__main__':
    cases = []
    for nstruct_i in range(1, nstruct + 1 ):
        for case_name in os.listdir('inputs'):
            case_path = os.path.join( 'inputs', case_name )
            for f in os.listdir(case_path):
                if f.endswith('.pdb'):
                    input_pdb_path = os.path.join( case_path, f )
                    break

            with open( os.path.join( case_path, 'chains_to_move.txt' ), 'r' ) as f:
                chains_to_move = f.readlines()[0].strip()

            cases.append( (case_name, case_path, input_pdb_path, chains_to_move, nstruct_i) )

    if use_multiprocessing:
        pool = multiprocessing.Pool( processes = min(max_cpus, multiprocessing.cpu_count()) )

    for args in cases:
        if use_multiprocessing:
            pool.apply_async( run_flex_ddg, args = args )
        else:
            run_flex_ddg( *args )

    if use_multiprocessing:
        pool.close()
        pool.join()
```
