## Supplementary material for "Distinct genetic pathways define pre-leukemic and compensatory clonal hematopoiesis in Shwachman-Diamond syndrome": Rosetta scripts: run.sbatch

```
#!/bin/bash
#SBATCH -c 12                               # Request one core
#SBATCH -N 1                               # Request one node (if you request more than one core with -c, also using
#SBATCH -t 0-12:00                         # Runtime in D-HH:MM format
#SBATCH -p short                           # Partition to run in
#SBATCH --mem=8G                           # Memory total in MB (for all cores)
#SBATCH -o runlogs/%j.out
#SBATCH -e runlogs/%j.err
#SBATCH --mail-type=END                    # Type of email notification- BEGIN,END,FAIL,ALL 
#SBATCH --mail-user=

module load gcc/6.2.0
module load python/2.7.12 
python eif6.py
```
