## Supplementary material for "Distinct genetic pathways define pre-leukemic and compensatory clonal hematopoiesis in Shwachman-Diamond syndrome": Rosetta scripts: EIF6_missense.sh

```
#!/bin/bash
sbatch  --export=RESNUM='1',MUTANT='T' run.sbatch; sleep 1
sbatch  --export=RESNUM='3',MUTANT='D' run.sbatch; sleep 1
sbatch  --export=RESNUM='5',MUTANT='D' run.sbatch; sleep 1
sbatch  --export=RESNUM='9',MUTANT='D' run.sbatch; sleep 1
sbatch  --export=RESNUM='12',MUTANT='K' run.sbatch; sleep 1
sbatch  --export=RESNUM='13',MUTANT='N' run.sbatch; sleep 1
sbatch  --export=RESNUM='14',MUTANT='D' run.sbatch; sleep 1
sbatch  --export=RESNUM='15',MUTANT='Y' run.sbatch; sleep 1
sbatch  --export=RESNUM='16',MUTANT='S' run.sbatch; sleep 1
sbatch  --export=RESNUM='17',MUTANT='P' run.sbatch; sleep 1
sbatch  --export=RESNUM='19',MUTANT='F' run.sbatch; sleep 1
sbatch  --export=RESNUM='20',MUTANT='I' run.sbatch; sleep 1
sbatch  --export=RESNUM='23',MUTANT='H' run.sbatch; sleep 1
sbatch  --export=RESNUM='23',MUTANT='N' run.sbatch; sleep 1
sbatch  --export=RESNUM='25',MUTANT='R' run.sbatch; sleep 1
sbatch  --export=RESNUM='27',MUTANT='V' run.sbatch; sleep 1
sbatch  --export=RESNUM='29',MUTANT='V' run.sbatch; sleep 1
sbatch  --export=RESNUM='36',MUTANT='I' run.sbatch; sleep 1
sbatch  --export=RESNUM='36',MUTANT='T' run.sbatch; sleep 1
sbatch  --export=RESNUM='46',MUTANT='S' run.sbatch; sleep 1
sbatch  --export=RESNUM='47',MUTANT='L' run.sbatch; sleep 1
sbatch  --export=RESNUM='51',MUTANT='P' run.sbatch; sleep 1
sbatch  --export=RESNUM='56',MUTANT='R' run.sbatch; sleep 1
sbatch  --export=RESNUM='56',MUTANT='Y' run.sbatch; sleep 1
sbatch  --export=RESNUM='57',MUTANT='P' run.sbatch; sleep 1
sbatch  --export=RESNUM='59',MUTANT='M' run.sbatch; sleep 1
sbatch  --export=RESNUM='61',MUTANT='H' run.sbatch; sleep 1
sbatch  --export=RESNUM='65',MUTANT='V' run.sbatch; sleep 1
sbatch  --export=RESNUM='66',MUTANT='S' run.sbatch; sleep 1
sbatch  --export=RESNUM='66',MUTANT='K' run.sbatch; sleep 1
sbatch  --export=RESNUM='67',MUTANT='W' run.sbatch; sleep 1
sbatch  --export=RESNUM='69',MUTANT='S' run.sbatch; sleep 1
sbatch  --export=RESNUM='69',MUTANT='V' run.sbatch; sleep 1
sbatch  --export=RESNUM='72',MUTANT='G' run.sbatch; sleep 1
sbatch  --export=RESNUM='73',MUTANT='R' run.sbatch; sleep 1
sbatch  --export=RESNUM='73',MUTANT='L' run.sbatch; sleep 1
sbatch  --export=RESNUM='78',MUTANT='Y' run.sbatch; sleep 1
sbatch  --export=RESNUM='80',MUTANT='K' run.sbatch; sleep 1
sbatch  --export=RESNUM='80',MUTANT='G' run.sbatch; sleep 1
sbatch  --export=RESNUM='81',MUTANT='R' run.sbatch; sleep 1
sbatch  --export=RESNUM='85',MUTANT='H' run.sbatch; sleep 1
sbatch  --export=RESNUM='92',MUTANT='E' run.sbatch; sleep 1
sbatch  --export=RESNUM='96',MUTANT='W' run.sbatch; sleep 1
sbatch  --export=RESNUM='99',MUTANT='Q' run.sbatch; sleep 1
sbatch  --export=RESNUM='99',MUTANT='K' run.sbatch; sleep 1
sbatch  --export=RESNUM='103',MUTANT='P' run.sbatch; sleep 1
sbatch  --export=RESNUM='105',MUTANT='D' run.sbatch; sleep 1
sbatch  --export=RESNUM='106',MUTANT='S' run.sbatch; sleep 1
sbatch  --export=RESNUM='111',MUTANT='H' run.sbatch; sleep 1
sbatch  --export=RESNUM='111',MUTANT='D' run.sbatch; sleep 1
sbatch  --export=RESNUM='111',MUTANT='S' run.sbatch; sleep 1
sbatch  --export=RESNUM='111',MUTANT='K' run.sbatch; sleep 1
sbatch  --export=RESNUM='112',MUTANT='N' run.sbatch; sleep 1
sbatch  --export=RESNUM='112',MUTANT='V' run.sbatch; sleep 1
sbatch  --export=RESNUM='113',MUTANT='C' run.sbatch; sleep 1
sbatch  --export=RESNUM='120',MUTANT='V' run.sbatch; sleep 1
sbatch  --export=RESNUM='122',MUTANT='V' run.sbatch; sleep 1
sbatch  --export=RESNUM='127',MUTANT='K' run.sbatch; sleep 1
sbatch  --export=RESNUM='129',MUTANT='Q' run.sbatch; sleep 1
sbatch  --export=RESNUM='133',MUTANT='P' run.sbatch; sleep 1
sbatch  --export=RESNUM='135',MUTANT='M' run.sbatch; sleep 1
sbatch  --export=RESNUM='149',MUTANT='E' run.sbatch; sleep 1
sbatch  --export=RESNUM='150',MUTANT='R' run.sbatch; sleep 1
sbatch  --export=RESNUM='158',MUTANT='R' run.sbatch; sleep 1
sbatch  --export=RESNUM='158',MUTANT='V' run.sbatch; sleep 1
sbatch  --export=RESNUM='159',MUTANT='R' run.sbatch; sleep 1
sbatch  --export=RESNUM='159',MUTANT='E' run.sbatch; sleep 1
sbatch  --export=RESNUM='160',MUTANT='P' run.sbatch; sleep 1
sbatch  --export=RESNUM='161',MUTANT='G' run.sbatch; sleep 1
sbatch  --export=RESNUM='167',MUTANT='N' run.sbatch; sleep 1
sbatch  --export=RESNUM='170',MUTANT='P' run.sbatch; sleep 1
sbatch  --export=RESNUM='173',MUTANT='P' run.sbatch; sleep 1
sbatch  --export=RESNUM='173',MUTANT='R' run.sbatch; sleep 1
sbatch  --export=RESNUM='177',MUTANT='R' run.sbatch; sleep 1
sbatch  --export=RESNUM='179',MUTANT='D' run.sbatch; sleep 1
sbatch  --export=RESNUM='179',MUTANT='G' run.sbatch; sleep 1
sbatch  --export=RESNUM='181',MUTANT='P' run.sbatch; sleep 1
sbatch  --export=RESNUM='181',MUTANT='H' run.sbatch; sleep 1
sbatch  --export=RESNUM='184',MUTANT='R' run.sbatch; sleep 1
sbatch  --export=RESNUM='184',MUTANT='E' run.sbatch; sleep 1
sbatch  --export=RESNUM='187',MUTANT='I' run.sbatch; sleep 1
sbatch  --export=RESNUM='187',MUTANT='K' run.sbatch; sleep 1
sbatch  --export=RESNUM='188',MUTANT='P' run.sbatch; sleep 1
sbatch  --export=RESNUM='194',MUTANT='T' run.sbatch; sleep 1
sbatch  --export=RESNUM='195',MUTANT='D' run.sbatch; sleep 1
sbatch  --export=RESNUM='196',MUTANT='R' run.sbatch; sleep 1
sbatch  --export=RESNUM='196',MUTANT='V' run.sbatch; sleep 1
sbatch  --export=RESNUM='196',MUTANT='E' run.sbatch; sleep 1
sbatch  --export=RESNUM='198',MUTANT='M' run.sbatch; sleep 1
sbatch  --export=RESNUM='200',MUTANT='I' run.sbatch; sleep 1
sbatch  --export=RESNUM='200',MUTANT='T' run.sbatch; sleep 1
sbatch  --export=RESNUM='204',MUTANT='D' run.sbatch; sleep 1
sbatch  --export=RESNUM='206',MUTANT='R' run.sbatch; sleep 1
sbatch  --export=RESNUM='206',MUTANT='Y' run.sbatch; sleep 1
sbatch  --export=RESNUM='207',MUTANT='R' run.sbatch; sleep 1
sbatch  --export=RESNUM='207',MUTANT='D' run.sbatch; sleep 1
sbatch  --export=RESNUM='210',MUTANT='I' run.sbatch; sleep 1
sbatch  --export=RESNUM='215',MUTANT='P' run.sbatch; sleep 1
```
