## Supplementary material for "Distinct genetic pathways define pre-leukemic and compensatory clonal hematopoiesis in Shwachman-Diamond syndrome": Rosetta scripts: run.sbatch

```
#!/bin/bash
#SBATCH -c 1                               # Request one core
#SBATCH -N 1                               # Request one node (if you request more than one core with -c, also using
#SBATCH -t 0-00:30                         # Runtime in D-HH:MM format
#SBATCH -p priority                        # Partition to run in
#SBATCH --mem=1G                           # Memory total in MB (for all cores)
#SBATCH -o runlogs/%j.out
#SBATCH -e runlogs/%j.err

DIRECTORY=/home/jwb29/eif6
PDB=/home/jwb29/eif6/prep_structure/output/full_model_18244_1_1992614_47_0001_0001.pdb
SCRIPT=${DIRECTORY}/eif6_mutants.xml
OUTPUT=${DIRECTORY}/output

/n/data1/bch/hemonc/bahl/samaneh/Rosetta/main/source/bin/rosetta_scripts.default.linuxgccrelease \
	-beta \
	-linmem_ig 10 \
	-in:file:s ${PDB} \
	-parser:protocol ${SCRIPT} \
	-parser:script_vars \
		resnum=${RESNUM}\
		mutant=${MUTANT}\
	-out:path:all ${OUTPUT} \
	-out:suffix _${RESNUM}${MUTANT} \
	-nstruct 1 \
```
